# Pectin induced colony expansion of soil-derived Flavobacterium strains

**DOI:** 10.1101/2020.06.26.174714

**Authors:** Judith Kraut-Cohen, Orr H. Shapiro, Barak Dror, Eddie Cytryn

## Abstract

The genus *Flavobacterium* is characterized by the capacity to metabolize complex organic compounds and a unique gliding motility mechanism. They are often abundant in root microbiomes of various plants, but the factors contributing to this high abundance are currently unknown. In this study, we evaluated the effect of various plant-associated poly- and mono-saccharides on colony expansion of two *Flavobacterium* strains. Both strains were able to spread on pectin and other polysaccharides such as microcrystalline cellulose. However, only pectin (but not pectin monomers), a component of plant cell walls, enhanced colony expansion on solid surfaces in a dose- and substrate-dependent manner. On pectin, flavobacteria exhibited bi-phasic motility, with an initial phase of rapid expansion, followed by growth within the colonized area. Proteomic and gene expression analyses revealed significant induction of carbohydrate metabolism related proteins when flavobacteria were grown on pectin, including selected SusC/D, TonB-dependent glycan transport operons. Our results show a positive correlation between colony expansion and the upregulation of proteins involved in sugar uptake, suggesting an unknown linkage between specific operons encoding for glycan uptake and metabolism and flavobacterial expansion. Furthermore, within the context of flavobacterial-plant interactions, they suggest that pectin may facilitate flavobacterial expansion on plant surfaces in addition to serving as an essential carbon source.

**Contribution to the field statement:** Members of the *Flavobacterium* genus are frequently rhizosphere competent, *i.e.* abundant and generally enriched on root surfaces relative to surrounding bulk soil, and previous studies suggest that they may play a role in plant health and ecosystem functioning. However, little is known about genetic and physiological factors that facilitate flavobacterial colonization and proliferation in this highly competitive environment.

In this study we found that pectin stimulates flavobacterial colonies in a bi-phasic manner, initially characterized by rapid expansion on agar followed by increased biomass production. This appears to be linked to pectin-stimulated induction of specific TonB-associated proteins evidentially involved in the binding and uptake of complex sugars. Given the fact that pectin and other glycans are a primary component of plant cell walls, we hypothesize that these mechanisms are at least partially responsible for the rhizosphere competence of flavobacteria.

## Introduction

The complex interactions between plant-associated microorganisms and their hosts (collectively referred to as the “plant holobiont”) are crucial for plant health and growth (Berendsen et al., 2012; Bulgarelli et al., 2013; Reinhold-Hurek et al., 2015). Plants modulate their rhizospheres, by exuding various small molecular weight compounds (rhizodeposits) such as sugars, amino acids and organic acids, by rhizodepositing root cap border cells, and by releasing various mono- and poly-saccharides in their mucilage (Dennis et al., 2010; Barret et al., 2011; Beauregard et al., 2013; Reinhold-Hurek et al., 2015; Massalha et al., 2017; Sasse et al., 2018). Collectively, these rhizodeposits, create a nutrient rich environment relative to the surrounding bulk soil, which facilitates colonization by soil microorganisms. Root colonizing bacteria can outcompete other soil bacteria by a combination of specific traits collectively coined “rhizosphere competence”, which include: motility, resistance to stress, ability to utilize plant-derived organic compounds, chemotaxis, and the production of secondary metabolites (Barret et al., 2011). Some earlier studies demonstrated that root-associated (rhizoplane) bacteria are attracted to the roots by plant-exuded organic acids such as malic, citric or fumaric acid and various amino acids (Rudrappa et al., 2008; Williams et al., 2008; Oku et al., 2012; Webb et al., 2017; Feng et al., 2019). Many of these sensed chemo-attractants, can also be consumed by the bacteria (Cremer et al., 2019). Although root recruitment and colonization mechanisms of certain plant-growthpromoting rhizobacteria (PGPR) have been identified and characterized (Lugtenberg and Dekkers, 1999; Yan et al., 2008; Pieterse et al., 2014), those responsible for recruitment of the vast majority of rhizoplane bacteria are currently an enigma.

*Flavobacterium* is a Gram-negative genus of bacteria from the phylum *Bacteroidetes* known to degrade complex organic compounds in freshwater, marine and soil environments (Kolton et al., 2013; McBride et al., 2014). It is highly abundant in the rhizoplane of a wide array of plants, in contrast to considerably lower abundance in the rhizosphere and bulk soil (Johansen et al., 2002; Janssen, 2006; Manter et al., 2010; Kolton et al., 2011, 2013; Lundberg et al., 2012; Bodenhausen et al., 2013; Bulgarelli et al., 2015). Several root- and soil-derived flavobacteria were found to antagonize various plant pathogens in different crops (Gunasinghe et al., 2004; Sang and Kim, 2012; Kolton et al., 2014a; Xue et al., 2015; Kwak et al., 2018), and recently it was discovered that endophytic *Flavobacterium* strains play a fundamental role in antagonizing phytopathogenic fungi, through specific secondary metabolites (Carrión et al., 2019). Furthermore, a recent study that applied bacterial network analysis indicated that Flavobacteria are potential drivers of pathogen suppression in root ecosystems (Wei et al., 2019). In addition, selected members of this genus have been characterized as plant growth promoting rhizobacteria (PGPR) of various crops (Hebbar et al., 1991; Alexander and Stewart, 2001; Gunasinghe et al., 2004; Manter et al., 2010; Sang et al., 2011; Sang and Kim, 2012).

Soil flavobacteria have specialized ability to decompose complex plant derived polysaccharides, such as pectin and glucomannan and the ability to secret various carbohydrate-active enzymes via the *Bacteroidetes* -specific type IX secretion system (T9SS) (McBride et al., 2009; Kolton et al., 2013; Kharade and McBride, 2015; McBride and Nakane, 2015; Lauber et al., 2018). Like other *Bacteroidetes,* they contain a myriad of genes that encode Polysaccharide Utilization Loci (PULs) that are activated specifically to facilitate glycan capture and sugar uptake (Martens et al., 2009; Jiménez et al., 2015; Foley et al., 2016). These PULs include outer membrane protein transducers involved in polysaccharide utilization, which are part of the TonB family, generally referred to as Starch Utilization System (SUS) proteins. Interestingly, comparative genomics revealed that genomes of soil and root-associated flavobacterial strains are significantly enriched with genes associated with plant polysaccharide degradation relative to aquatic strains from this genus, indicating that physiology of this genus is strongly influenced by its ecology (Kolton et al., 2013).

Most terrestrial *Flavobacterium* strains possess a unique gliding mechanism that rapidly propels them over solid and semi-solid surfaces. The proteins that comprise this gliding system are molecularly intertwined with at least fifteen proteins that make up the T9SS, seven of whom are responsible for the secretion of SprB and RemA adhesins which are expressed on the cell surface and involved in gliding (Shrivastava et al., 2013; Johnston et al., 2018). We previously demonstrated that this T9SS-gliding complex is crucial for root colonization by flavobacteria, and this colonization was positively linked to the induction of plant resistance to foliar pathogens (Kolton et al., 2014a).

Collectively, the above studies strongly suggest that terrestrial flavobacterial strains have evolved means that enable them to interact with plant roots, and that these interactions are beneficial to plant health. Nonetheless, the specific mechanisms behind this phenomenon are currently unclear. In this study, we assessed the impact of an array of plant cell wall-derived substrates on the motility and growth dynamics of flavobacteria by coupling conventional petri dish assays and live-imaging fluorescent microscopy with proteomic and gene expression analyses. We demonstrate that pectin, a plant cell wall-associated polysaccharide, facilitates bi-phasic proliferation over solid surfaces through induction of specific TonB-associated glycan uptake operons. These results suggests that the link between pectin, motility and carbohydrate metabolism may be fundamental to rhizosphere competence in flavobacteria.

## Materials and methods

### Bacterial Strains and growth conditions

Flavobacterial strains targeted in this study included *F. johnsoniae* strain UW101 (ATCC17061), a gliding/secretion impaired *(ΔgldJ F. johnsoniae* strain UW101(Braun and McBride, 2005), *Flavobacterium* sp. F52, isolated from the roots of a greenhouse pepper (Sela et al., 2012; Kolton et al., 2014b), and a gliding/secretion impaired (Δ*gldJ Flavobacterium* sp. F52 strain (Kolton et al., 2014b). For live imaging microscopy (see below) the fluorescent strain *F. johnsoniae* UW101 WT+pAS43 (Flavo GFP) Flavo-ErytRFlavo-CefR, and the gliding/secretion impaired *F. johnsoniae* UW102-48 *(ΔgldJ+pAS43* (Flavo GFP) Flavo-ErytR Flavo-CefR were used (Mcbride and Baker, 1996; Staroscik et al., 2008). Erythromycin, 100 ug/ml was added to the media of the GFP labeled bacteria.

Flavobacteria strains were grown in CYE medium (Casitone, 10 mg/ml, yeast extract at 5mg/ml, and 8 mM MgSO4 in 10 mM Tris buffer [pH 7.6]) at 30°C, as previously described (Mcbride and Baker, 1996). To observe colony-spreading, bacteria were grown on PY2 agar medium (2 g peptone, 0.5 g yeast extract,10 mg/ml agar, pH 7.3) at 30°C (Agarwal S. et al., 2001).

### Organic amendments to growth media

A suite of mono- and polysaccharides were amended to growth media in various configurations (as described below), to evaluate colony spreading dynamics of the selected flavobacterial strains. The following substances were used in this study: pectin (Sigma P9135), D(+)glucose (Fischer scientific 10373242), microcrystalline cellulose (M.cellulose-partially depolymerized cellulose synthesized from an α-cellulose precursor, Merck 102331), D(-)arbinose (Acros 161451000), glucomannan (Megazyme P-GLCML), L-rhamnose (Sigma 83650) and D(+)-galacturonic acid monohydrate (Sigma 48280-5G-F), Polyethylene Glycol (PEG) 8000 (Amresco 0159). All substances were dissolved and suspended to 2% final concentration in double distilled water (DDW), unless indicated otherwise. Glucose, arabinose, rhamnose and galacturonic acid (titrated to pH 7) were dissolved and filtered through a 0.22-micron filter. Pectin was dissolved in DDW heated at 80°C, and subsequently filtered through 0.45-micron filters. M.cellulose, and glucomannan were mixed with DDW heated to 50°C and then autoclaved for 20 min.

### Flavobacterial growth on various plant-derived poly- and mono-saccharides

The selected plant-derived poly- and mono-saccharides were amended to the PY2 agar plates in two manners: (i) to test the effect of specific carbon sources on the directional proliferation of flavobacteria, 10 μl of the selected compounds were thinly applied along a line projecting outward from the center of the petri dish using a pipetor (**Fig S1**); (ii) to test the effect of specific carbon sources on the general proliferation of flavobacteria, 500μl from each 2% sugar solution were uniformly smeared over the entire petri dish (**Fig 1A, C, Fig 2A**). Non-soluble substances such as M.cellulose were vigorously vortexed and dispensed using a cut tip. Where two sugars were used (*i.e.* rhamnose+ galacturonic acid) we added 250μl of each. In all cases, plates were left to dry overnight after adding organic amendments. Flavobacteria were incubated on CYE media overnight, colonies were harvested and subsequently diluted in 200ul saline to 0.6-1 OD (5×10^9^-1.12 X10^10^ cells), vortexed well, and 2ul were spotted in the center of PY2 agar covered or streaked with the selected plant-derived poly- and monosaccharides as indicated above. Plates were left to dry for 15min and then incubated for 48 hours at 30°C. The colony area or the length of expansion (in cm) were measured using Fiji (Schindelin et al., 2012) and statistics was calculated using JMP®, Version Pro 14 (SAS Institute Inc., Cary, NC, 1989-2019). Differences between length/area were considered as significantly different when p<0.05 in Tukey HSD test unless indicated differently.

**Fig 1.**
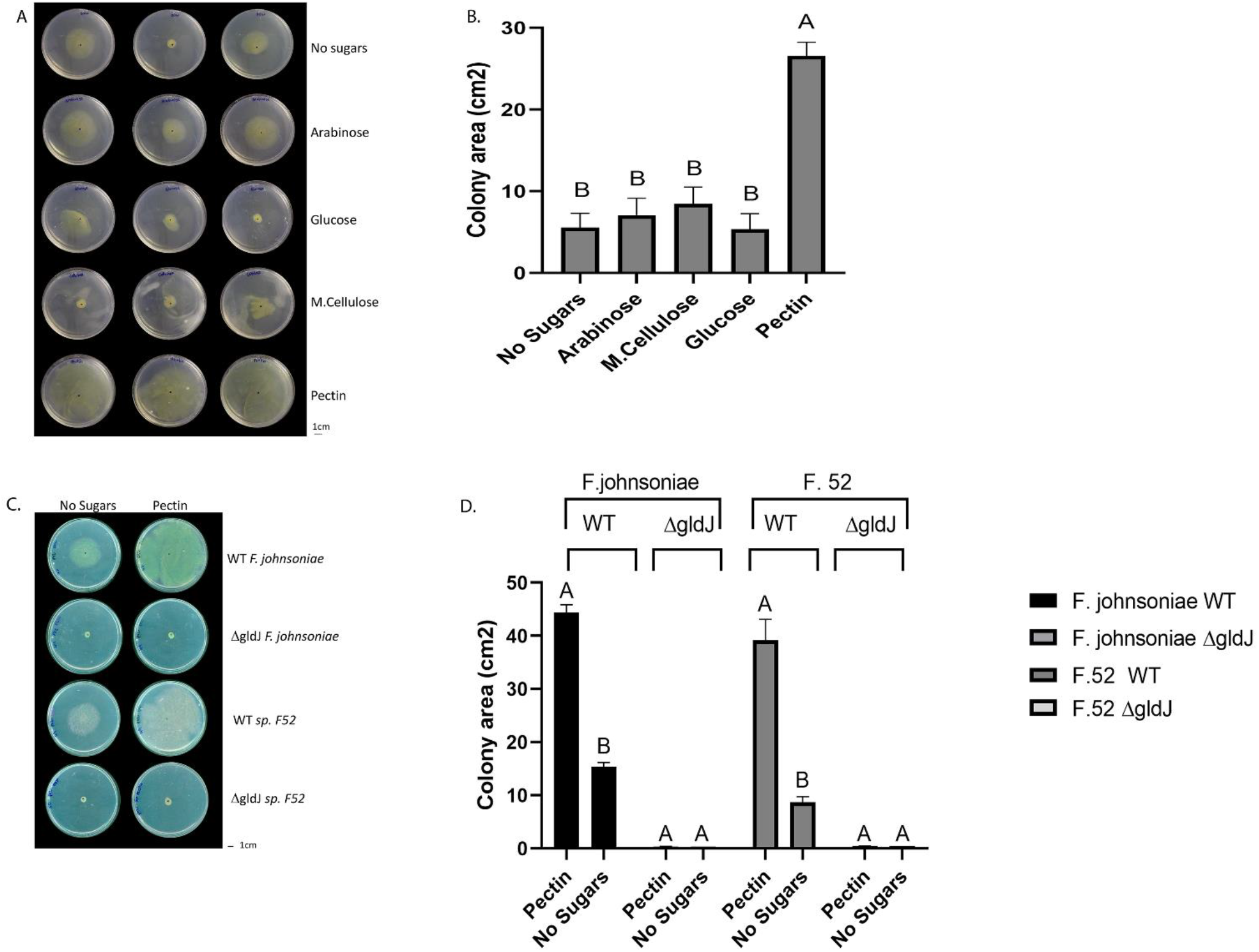
Impact of plant-derived poly- and mono- saccharides on proliferation of flavobacteria. Image (A) and quantification (B) of colony area of *F. johnsoniae* colony expansion on PY2 agar amended with different mono- and polysaccharides: arabinose, glucose, M.cellulose (M.Cellulose), pectin and the no sugar (DDW control), after 48hr incubation at 30°C (N=4). Figure represents one experiment (out of four) with 3 technical repeats in each. Error bars represents standard error. Image (C) and quantification (D) of colony expansion on PY2 agar of wild type (WT) and gliding mutants *(ΔgldJ)* of *F. johnsoniae* and *Flavobacterium* sp. F52, amended with DDW (no sugar) or 2% pectin after 48hr incubation at 30°C (N=3). Colony area was measured using Fiji. Letters indicate statistical significance calculated using ®JMP Pro14, was considered significant if p<0.05 by Tukey HSD. Error bars represents standard error.

**Fig 2.**
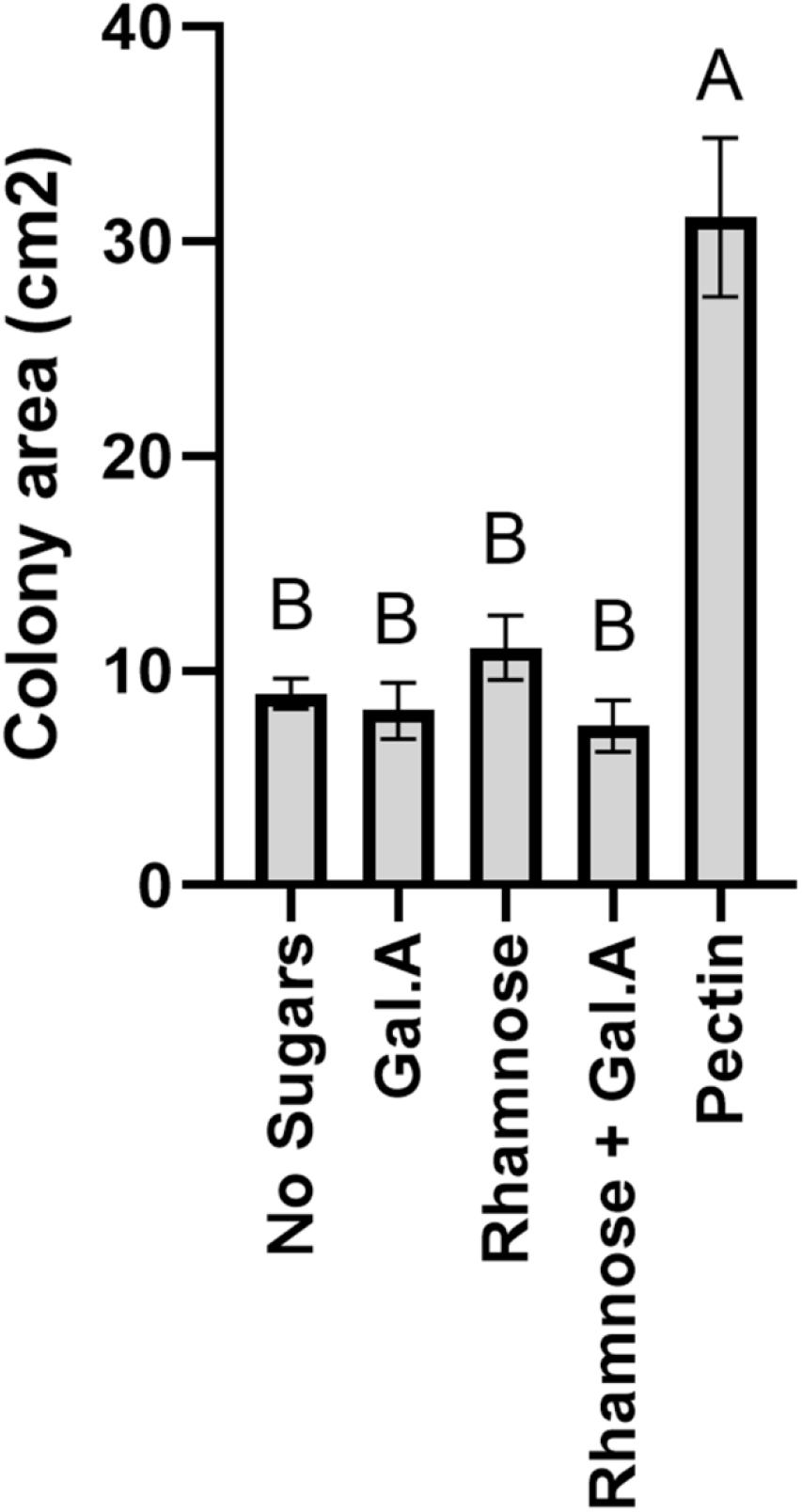
Impact of pectin precursors (galacturonic acid and rhamnose) on proliferation of flavobacteria. Graphic description of *F. johnsoniae* colony area on PY2 agar amended with DDW, pectin galacturonic acid, rhamnose and galacturonic acid and rhamnose incubated at 30°C for 48 h (N=8). Statistics significance was calculated using ®JMP Pro14, and means was considered significant when p<0.05 by Tukey HSD. Error bars represents standard error.

### Live Imaging Fluorescent Microscopy Experiments

To visualize the effect of different mono- and polysaccharides on *F. johnsoniae* colony expansion, we filled the wells of a 24 well plate with 500μl of PY2 agar. Subsequently, 10ul of the mono- and polysaccharides (2%) or the control (DDW) were gently applied to the wells in triplicates, plates were rotated for 1hour and dried overnight at room temperature. A 30G needle (KDL 16-0310) was used to seed the bacteria in the center of each well by punching an agar plate with a 24hr grown colony. Microscopic imaging was performed using a NIKON eclipse Ti microscope (Nikon, Japan) equipped with a ProScan motorized XY stage, an HF110A system (enabling rapid switching of emission filters)(Prior Scientific, MA, USA) and a temperature-controlled stage incubator (on 25°C) (LAUDA ECO RE 415, Korea). Bright field illumination was provided by a cool LED pE-100A (Cool LED, UK). Excitation light for epifluorescence microscopy was provided by a Spectra X light engine (Lumencor, USA). Imaging was performed using a long working distance 40X objective (NA 0.6) (Nikon, Japan). Images were captured at 2 hr intervals for 44 hr using an ANDOR zyla 5.5 MP sCMOS camera (Oxford Instruments, UK).

Images were processed using the NIS elements AR 4.6 (64 bit) software package (Nikon, Japan) and Fiji (Schindelin et al., 2012). Fluorescence level was normalized to the initial measured value (to avoid differences in the initial number of seeded bacteria) and to the maximal fluorescence on PY2 media amended with DDW (to reduce variability of GFP fluorescence levels between movies). The population growth dynamic of GFP-*F. johnsoniae* on each substance computed using Fiji’s time series analyzer plugin, and average fluorescence density profiles of the expanding population over time was quantified using JMP®, Version Pro 14. (SAS Institute Inc., Cary, NC, 1989-2019).

### Proteomic Sample preparation

Wild type *F. johnsoniae,* was grown on PY2 agar plates smeared either with 500μl of 2% pectin or with DDW, in triplicates. After 48hr, bacteria were scraped from plates in 1ml of 4°C PBS and centrifuged for 10 min, 4500rpm in 4°C. Supernatant was discarded and bacterial pellets were processed.

Samples were subjected to in-solution tryptic digestion using a modified Filter Aided Sample Preparation protocol (FASP). Sodium dodecyl sulfate buffer (SDS) included: 4% (w/v) SDS, 100mM Tris/HCl pH 7.6, 0.1M DTT. Urea buffer (UB): 8 M urea (Sigma, U5128) in 0.1 M Tris/HCl pH 8.0 and 50mM Ammonium Bicarbonate. Bacterial cell pellets were lysed in SDS buffer. Lysate was centrifuged at 16,000 g for 10min. 50 ug total protein were mixed with 2 mL UB, loaded onto 30 kDa molecular weight cutoff filters (Sartorius VS15RH22) and centrifuged. 1.5 ml of UB were added to the filter unit, and centrifuged at 14,000 g for 40 min. Proteins were alkylated using iodoacetamide (10 mM final concentration) and washed twice with Ammonium Bicarbonate. Trypsin was then added (50:1 protein amount:trypsin) and samples incubated at 37°C overnight, followed by a second trypsin digestion for 4 hours at 37°C. Digested peptides were then collected in a clean tube by centrifugation, acidified with trifloroacetic acid, desalted using HBL Oasis (Waters 094225), speed vacuumed to dryness and stored in −80°C until analysis. All chemicals used were from Sigma Aldrich, unless stated otherwise. Sample preparation and initial statistical analysis was performed at the Nancy and Stephen Grand Israel National Center for Personalized Medicine, The Weizmann Institute of Science, Rehovot, Israel.

### Liquid chromatography and proteomics analysis

Each sample was loaded and analyzed using split-less nano-Ultra Performance Liquid Chromatography (10 kpsi nanoAcquity; Waters, Milford, MA, USA). The mobile phase was: A) H2O + 0.1% formic acid and B) acetonitrile + 0.1% formic acid. Desalting of the samples was performed online using a Symmetry C18 reversed-phase trapping column (180 μm internal diameter, 20 mm length, 5 μm particle size; Waters). The peptides were then separated using a T3 HSS nano-column (75 μm internal diameter, 250 mm length, 1.8 μm particle size; Waters) at 0.35 μL/min. Peptides were eluted from the column into the mass spectrometer using the following gradient: 4% to 20%B in 155 min, 20% to 90%B in 5 min, maintained at 90% for 5 min and then back to initial conditions. ULC/MS grade solvents were used for all chromatographic steps. Each sample was analyzed on the instrument separately in a random order in discovery mode. Raw data was searched against the *F. johnsoniae* protein databases, to which a list of common lab contaminants was added. Database search was done using the Mascot algorithm, and quantitative analysis was performed using the Expressionist software from GeneData. A student’s t-test based on log-transformed intensity values was performed determine significant differences in protein expression between colonies grown with and without pectin. As a rule of thumb we consider significant differences to be >1 peptide per protein, fold change >2 or <0.5, and <0.05 p-value. Proteins were functionally annotated using RAST (Rapid Annotations using Subsystems Technology) (Overbeek et al., 2014).

Twenty-five pectin-induced proteins were selected based on high fold change between PY2 agar coated with pectin vs. PY2 agar coated with DDW and statistical significance (proteins without annotation were removed from this analysis). Gene abundance across the dataset was normalized to 100% for each gene, and a heatmap was created using studio.plot.ly (https://plotly.com/).

The mass spectrometry proteomic data was deposited to the ProteomeXchange Consortium through the PRIDE partner repository (Perez-Riverol et al., 2019) under the dataset identifier PXD023649.

### RNA extraction

To assess the expression of selected genes in the presence of pectin, total RNA was extracted from *F. johnsoniae* cells grown for 48 hours at 30°C on PY2 plates covered with 500μl of DDW or 2% pectin as described above. For each plate, bacteria were suspended in 1 ml cold (4°C) PBS buffer, washed once in cold PBS, centrifuged for 2min at 18,000g, resuspended in TE buffer supplemented with 0.4mg/ml lysozyme and then incubated for 10 min at room temperature. RNA was subsequently extracted from cells using the TRIzol reagent (TRIzol® InvitrogenTM, #15596026), following the manufacturer’s instructions. Residual DNA was removed from the RNA samples by digesting with RQ1 DNAse (Promega M6101A) at 37°C for 40 min. For the real time experiments, cDNA was synthesized using 50ng of DNAse treated RNA, with 1ul of random primers (Promega C118A). Synthesis of single strand cDNA was achieved using ImProm-IITM Reverse-Transcriptase (Promega, Madison, WI, United States).

The integrity and concentration of the extracted RNA and cDNA, was examined with a QubitTM 3.0 Fluorometer (Thermo Fisher Scientific, United States) using reagents and protocols supplied by the manufacturer, and by electrophoresis of samples on a 0.8% agarose gel.

### Quantitative PCR Assessment of Gene Expression Levels

The expression of 10 genes encoding for proteins found to be significantly induced on pectin in the proteomic analyses were analyzed using qPCR. Primers for qRT-PCR experiments (**Table S5**) were constructed based on the *F. johnsoniae* genome sequence and were pre-designed using the PrimerQuest® tool (Integrated DNA Technologies, USA). Triplicate cDNA samples for each of the treatments (with or without pectin) were diluted and 2ng was used in a 20 μl final reaction volume together with 10ul Fast SYBR^TM^ green PCR master mix (Thermo Fisher scientific), 100 nM each of forward and reverse primers, DDW and 1ul of template cDNA. Amplification was carried out on a StepOnePlus real-time PCR thermocycler (Applied Biosystems, Foster City, CA, United States) using the following program: heat activation of DNA polymerase at 95°C for 3 min and 40 cycles at 95°C for 5 sec for denaturation and primer annealing and extension at 60°C for 30 sec. A melting curve was produced to confirm a single gene-specific peak and to detect primer-dimer formation by heating the samples from 60 to 95°C in 0.3°C increments. Gene amplification was confirmed by detection of a single peak in the melting curve analysis. For each gene, PCR gene amplification was carried out using three independent biological replicates. Expression of each of the targeted genes was normalized to that of three alternative housekeeping genes (16S rRNA, DNA gyrase subunit B *(gyrB,* EC 5.99.1.3), and the electron transfer flavoprotein, alpha subunit (ETF) threonine synthase (EC 4.2.3.1). These genes were selected because there was no detected difference in their expression when grown on PY2 media amended with pectin *vs.* DDW in the proteomic analysis. The relative abundance of each target gene relative to a reference gene was determined according to the method described previously (Livak and Schmittgen2001). Concentrations and ΔΔCT values were calculated and analyzed with the StepOne software v2.3 (Applied Biosystems, Foster City, CA, United States). Concomitant “no-RT” reactions, lacking reverse transcriptase, were performed for each sample and run to confirm absence of DNA contamination, as well as no template controls (NTCs) to confirm lack of contamination. Reaction efficiency was monitored by means of an in internal standard curve using a 10-fold dilution of DNA ranging from 0.01-10ng of DNA per reaction, in triplicates. Efficiency was between 92.1 and 97.2% for all primers, and R2-values were greater than 0.99. Data analysis was conducted using the StepOne software v2.3 (Applied Biosystems, Foster City, CA, United States).

## Results

### Growth of *Flavobacterium* strains on various carbon sources

We evaluated growth dynamics of flavobacteria on rich media (PY2 agar) coated with selected plant-derived poly- and mono-saccharides (**Fig 1A**). Colony expansion of *F. johnsoniae* on PY2 agar media coated with pectin was close to five times higher than on the same media coated with other analyzed mono- and polysaccharides or with DDW, (control) (p<0.05, Tukey-Kramer HSD test) (**Fig 1B**). Wild type (WT) and gliding/typeIX secretion system mutants *(ΔgldJ)* of *F. johnsoniae* and the pepper root isolate *Flavobacterium* sp. F52 were inoculated in the center of PY2 agar media amended with or without pectin (**Fig 1C**). When grown on pectin, WT colonies of both *Flavobacterium* strains significantly expanded after 48 hours of incubation, while growth was reduced in the control without pectin (DDW). In contrast, gliding mutant *(ΔgldJ)* colonies of both flavobacterial strains did not expand (**Fig 1C, D**), indicating that the gliding apparatus is a prerequisite for pectin-induced colony expansion.

### Dose-dependent pectin facilitated colony expansion

To determine whether expansion on pectin is dose dependent, *F. johnsoniae* and *Flavobacterium* sp.F52 strains were inoculated at the center of PY2 agar media plates streaked with pectin at final concentrations of 0.5, 1, 2 and 4%. For all the examined pectin concentrations colonies radiated along the pectin streaks, but expansion was more significant on 2% and 4% pectin (p<0.05,Tukey-Kramer HSD test) (**FigS1**).

Since galacturonic acid and rhamnose are the two major components of pectins, we examined the colony expansion of *F. johnsoniae* on PY2 agar coated with 2% titrated galacturonic acid, 2% rhamnose, a combination of galacturonic acid and rhamnose, 2% pectin and DDW. *F. johnsoniae* expansion on galacturonic acid, rhamnose (alone or combined) did not facilitate significant colony expansion (p<0.05,Tukey-Kramer HSD test) in contrast to colonies that were grown on pectin (**Fig 2**).

### Temporal dynamics of pectin induced *F. johnsoniae* colony expansion

The expansion of green fluorescent protein (GFP)-labeled *F. johnsoniae* on PY2 agar coated with glucose, M.cellulose, glucomannan, Peg8000, pectin or DDW (without sugar amendment), was visualized at a higher resolution using time-lapse microscopy. Colony morphology after 32 hours, on each tested substance is presented in **Fig 3A**.

**Fig 3.**
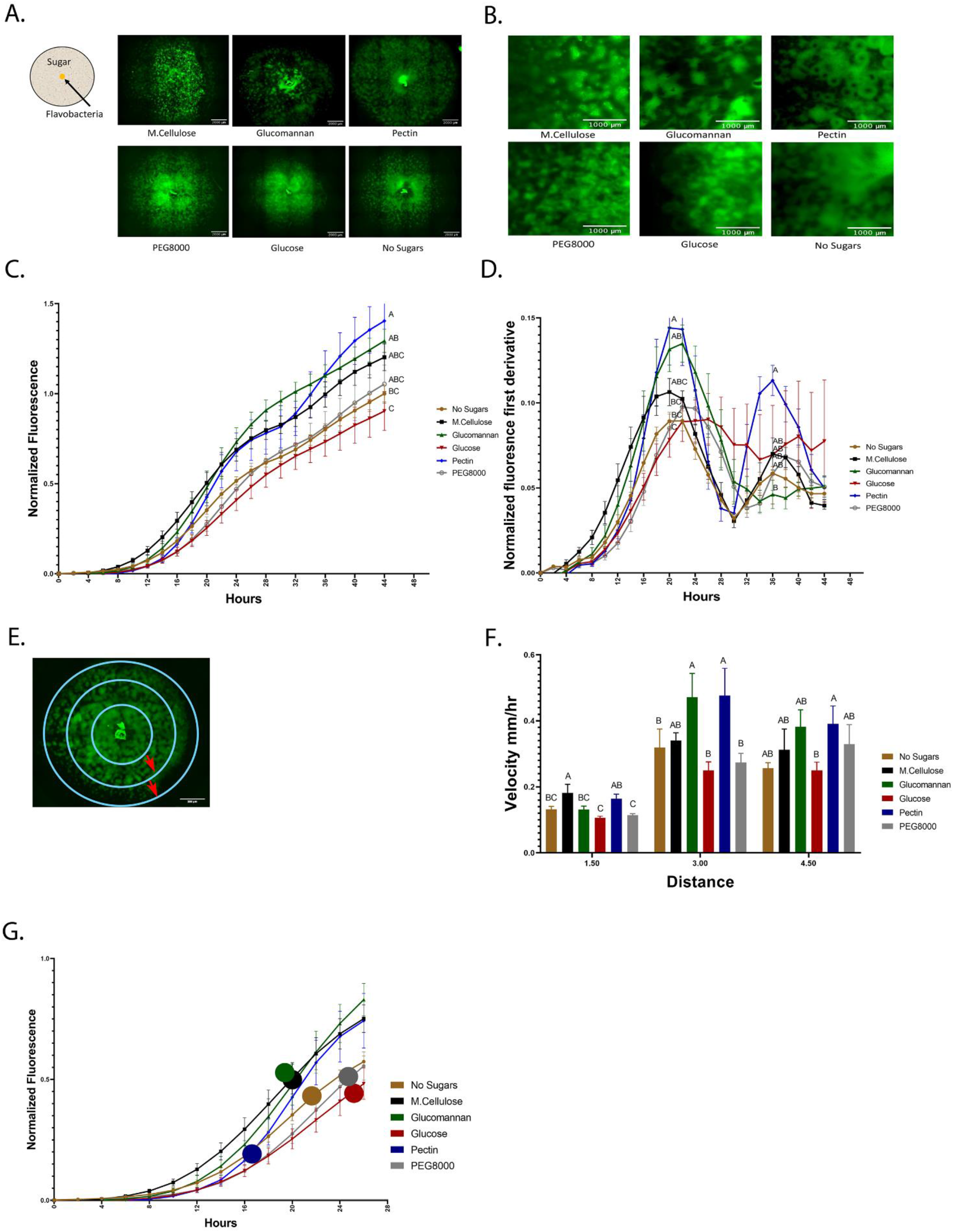
Temporal dynamics of flavobacterial proliferation on polysaccharides using live imaging microscopy. (A) Morphology of GFP-labeled *F. johnsoniae* colonies on PY2 agar amended with selected plant-derived poly- and mono-saccharides: M.cellulose, glucomannan, pectin, glucose and PEG8000 and DDW (no sugar control). Bacteria were inoculated in the center of PY2 agar coated with the indicated substances (schematically described in the insert). Images show colony morphology after 32hr. (B) Enlarged image of GFP*-F. johnsoniae* colony morphology after 16hr of growth on the selected plant-derived poly- and mono-saccharides as indicated in (A). (C) Growth rates of GFP*-F. johnsoniae* colonies on the selected plant-derived poly- and monosaccharides. Data was normalized as described in the materials and methods section. Differences in the average colony fluorescence intensity after 44 hr was compared and considered significant if p<0.05 by Tukey HSD (indicated by letters). Data includes means and data from three biological replicates composed of three technical repeats in each. (D) Temporal dynamics of GFP*-F. johnsoniae* growth rates. Growth was compared at the peaks (20hr and 36hr) and considered significant if p<0.05 by Tukey HSD. (E) Schematic diagram showing the three characterized regions of interest (ROI-1.5, 3 and 4.5 mm radii) used to evaluate of bacterial expansion rates. (F) Estimated expansion rates of GFP*-F. johnsoniae* on the selected plant-derived poly- and monosaccharides. Velocity was estimated by expansion time in hours taken to cross known ROIs as indicated in E. Differences between treatments were considered significant when p<0.05 by Student T test (G) Estimated expansion time relative to estimated growth of GFP*-F. johnsoniae* on the selected plant-derived poly- and monosaccharides. Colored circles mark time (h) for bacteria to cross the 4.5mm radius on each substance as calculated in F. All error bars represents standard error.

Growth dynamics were clearly affected by the sugar type. Expansion on pectin was characterized by a relatively long lag phase. However following this stage, colonies rapidly expanded and after 22 hours, the fluorescent signal of colonies grown on pectin surpassed both the control (without sugar amendment) and the glucose (according to Student T test test) and other amended compounds, after 36 hours. Plates coated with polyethylene glycol (PEG8000) served as an additional control to rule out mechanosensitivity associated with substrate stiffness (Tchoufag et al., 2019), since its viscosity is similar to that of pectin. Colony expansion on Peg8000 was similar to the DDW control without amended sugars, suggesting that effects were specific to pectin. Growth on pectin facilitated multiple ring-like microstructures that resemble previously described “raft” structures (Gorski et al., 1993) (**Fig 3B, Supplementary movie S1**). Conversely, expansion of colonies on glucose did not display this bi-phasic expansion. While bi-phasic growth was observed in all of the non-amended and amended plates, this phenomenon was most pronounced in the pectin-amended plates (**Supplementary movie S1**). Bacterial colonies grown on glucose expanded the least resulting in compact, small colonies with less ringed structures, suggesting that glucose has an inhibitory effect on *F. johnsoniae* motility and possibly growth as was previously demonstrated (**Fig 3C**) (Wolkin and Pate, 1984; Imamura et al., 2018). The most significant expansion observed between 6 to 20 hours was on M.cellulose, but the colonies grown on glucomannan and pectin proliferated at later times, and the colonies grown on pectin reached the greatest intensity at 44 hours (**Fig 3C**), thereby surpassing the colonies grown on other sugar sources (p<0.05,Tukey-Kramer HSD test). The high sensitivity of the fluorescence detection enabled us to visualize lateral colony expansion on M.cellulose, which was less visible on agar plates (**Fig 1A, movie S1).**

We further evaluated the pectin-induced bi-phasic expansion described above. An initial peak in fluorescence occurred at 20 hours, and a second peak at 36 hours (**Fig 3D**). After 20 hours of growth, total fluorescence was highest in cells grown on pectin, glucomannan and M.cellulose, and lowest on glucose (p<0.05, Tukey-Kramer HSD test).

Next, we estimated the velocity of colony expansion on the selected mono- and polysaccharides by measuring the time it took the colonies to cross three radials (3mm, 6mm and 9mm), and subsequently calculating the mean velocity from circle to circle (**Fig 3E**). In the first 1.5mm radius, the estimated colony expansion velocity was higher in pectin and M.cellulose and similar on the rest of the substances. Between 1.5-3mm, colony expansion on pectin and glucomannan increased. Colony expansion velocity in the outer circle was similar for all substance except glucose, with pectin still exhibiting the fastest expansion rates of all the tested mono- and polysaccharides although this was not statistically significant (**Fig 3F**). Thus, *F. johnsoniae* expanded faster on pectin than on any of the other tested carbon sources (**Fig 3G**). Collectively, we conclude that the bi-phasic growth of *F. johnsoniae* is very prominent on pectin. In summary, after an initial lag phase, *F. johnsoniae* is characterized by a rapid expansion phase, followed by a slower growth phase where cells appear to spread less and gain biomass.

### Specific TonB/Sus transducers are expressed in response to growth on pectin

In order to gain greater insight into the molecular mechanisms associated with flavobacterial colony expansion on pectin, we conducted a proteomic assay in which we examined differential intracellular protein expression after 44 hr growth of *F. johnsoniae* on PY2 agar coated with pectin relative to the same rich media coated with DDW. Eighty-three proteins were more expressed on pectin, whereas forty-three were more expressed on the DDW control (**Table S1A+B**). A substantial proportion of these proteins were unassigned, while a large fraction of the proteins (17%, 22% and 37% of KEGG, SEED and EggNog annotations, respectively) were associated with carbohydrate metabolism (**FigS2A-C**). Of the 25 most markedly pectin-induced proteins identified, 13 were involved in polysaccharide uptake, processing and metabolism, including four Sus C/D related proteins (**Fig 4A**). Other pectin-induced proteins included a novel transcriptional regulator (12-fold higher on pectin) and a protein associated with auxin regulation (26-fold higher on pectin) (**Table S1A+B**). Interestingly, none of the differentially synthesized proteins were gliding related (**Table S2**).

**Fig 4.**
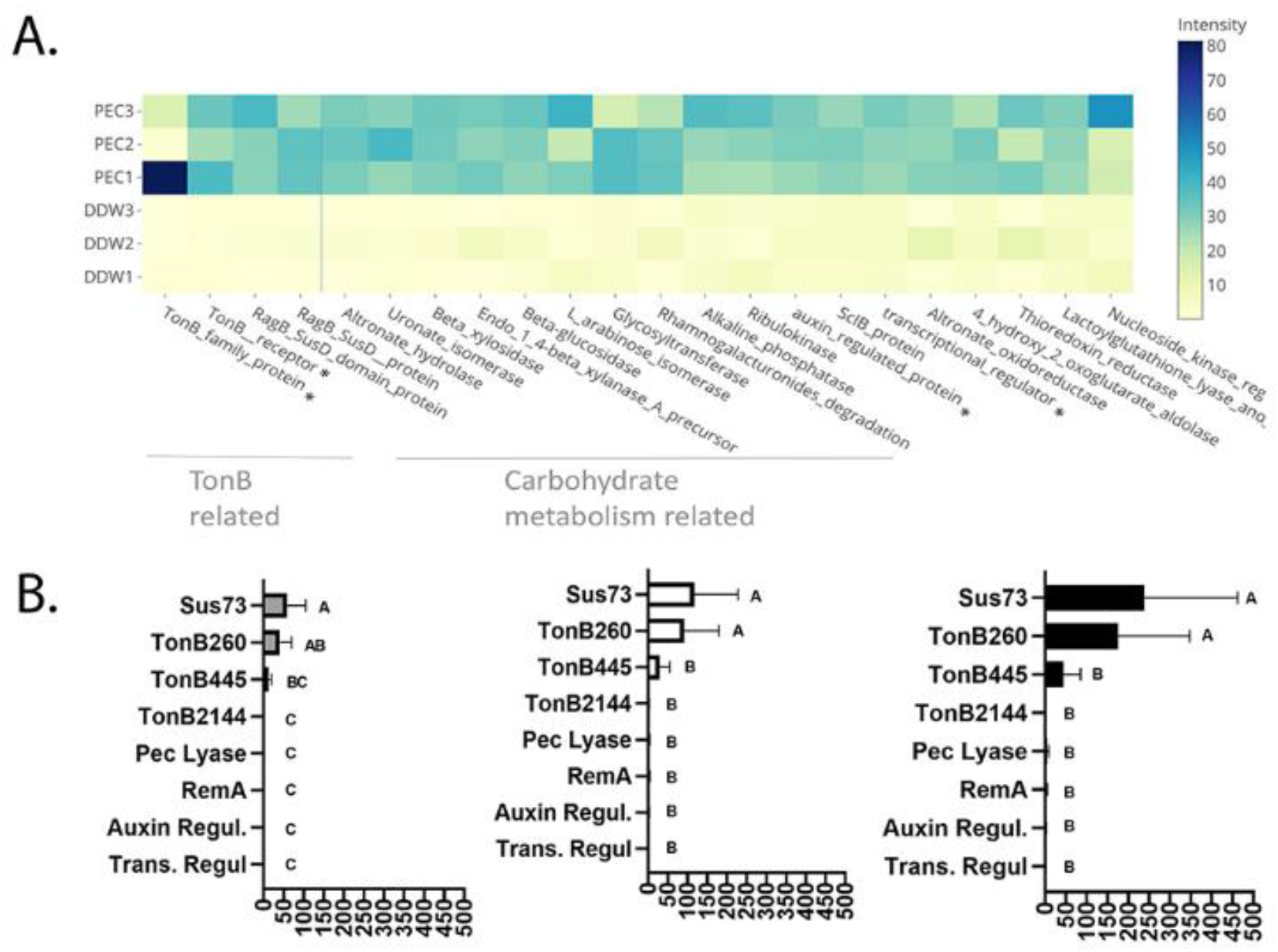
Pectin induced flavobacterial genes and proteins. (A) Differential expression of the 25 most substantial pectin induced proteins based on proteomic analysis of *F. johnsoniae* colonies grown on PY2 medium amended with pectin relative to colonies grown on identical media amended without pectin (DDW). Heat map shows triplicates for each treatment. All described proteins are statistically significant (p<0.05). The asterisk (*) marks proteins that were examined by qPCR. From left to right: TonB450, TonB260, Auxin regulator, Transcriptional regulator (B) The mRNA expression level of selected proteins Sus73 (SHH12854.1), TonB260 (WP_012026229.1), TonB445 (WP_012022294.1), TonB2144 (WP_012026069.1), Auxin regulator (WP_012023111.1), Transcriptional regulator (WP_012022876.1), Pectate lyase (WP_012026072.1) and RemA (WP_012022896.1) shown to be induced in the proteomic analysis described in (A), using quantitative real-time PCR (qPCR). Fold changes in mRNA levels of the target genes were normalized against the 16SrRNA gene (left), the Electron transfer Flavoprotein, alpha subunit (WP_012023552.1) (center), and the DNA gyrase subunit B (WP_012024321.1)(right). Change in target genes fold change RNA expression was calculated using the 2^-ΔΔCT^ method and statistical significance (p<0.05) by Student T-test. Error bars represent standard errors of six independent experiments based on two independent RNA extractions.

Of the 44 previously-described SusC and 42 SusD homologues identified in the *F. johnsoniae* genome (McBride et al., 2009), 27 SusC and 15 SusD proteins were detected in the proteomic analysis, in addition to 610 proteins encoding flanking genes surrounding these Sus proteins encoding genes that constitute PUL clusters seemingly associated with glycan metabolism. Of these, three SusC and six SusD proteins along with 31 associated PUL genes encoding proteins (forming 4 gene clusters **Table S3**) were significantly induced in response to growth on pectin in the proteomic analysis (**Table S4**).

In order to validate the proteomic results, *F. johnsoniae* cells were again grown on PY2 agar coated with pectin or DDW as described in the proteomic analysis, and the expression of 8 genes (**Table S5**) were evaluated by quantitative real time PCR. Pectin did not induce expression of *remA* (WP_012022896.1) encoding for an adhesion protein involved in gliding motility (not evaluated in the proteomic analysis since it’s an extra-membrane protein), indicating that synthesis of this protein is not enhanced on pectin at the tested time point. Expression of genes encoding for the novel transcriptional regulator (Trans. Regul WP_012022876.1), pectate lyase (Pec lyase WP_012026072.1), putative auxin-regulated annotated protein (Auxin WP_012023111.1) and tonB 2144 (WP_012026069.1), were also not differentially expressed on pectin-coated media despite the fact that they were significantly induced on pectin in the proteomic analysis. Among the examined TonB/SusC related genes, tonB 260 (WP_012026229.1) and sus73 (SHH12854.1) were significantly upregulated (60-100 fold, p<0.05 Tukey-Kramer HSD test) while TonB 445 (WP_012022294.1) was substantially upregulated (10-20 fold) but not statistically significant (**Fig 4B**). tonB260 and sus73 are part of an 18-gene cluster, of which all of the encoded proteins were upregulated in the presence of pectin in the proteomic analysis (**cluster 1-Table S3**). Using two different prediction tools, the TonB260 and Sus73 encoding genes were mapped to the same operon together with a gene encoding for the hydrolytic enzyme polygalacturonase that cleaves the α (1-4) bonds between adjacent galacturonic acids within the homogalacturonic acid backbone of pectin (**Table S6**).

## Discussion

### Pectin facilitates rapid spreading of flavobacteria

Results from this study demonstrated that flavobacteria not only metabolize complex plant cell wall-derived glycans but that these polysaccharides (particularly pectin) facilitate rapid spreading of flavobacteria over solid surfaces, even when carbon was not limited. Both the pattern and the extent of *F. johnsoniae* colony expansion was dictated by the carbon source supplemented to the growth media. The expansion on rich media coated with pectin and glucomannan was rapid, patchy, non-uniform and sparse relative to colonies grown on rich media coated with glucose or DDW. On a microscopic level, this rapid expansion was characterized by ring like micro-structures, resembling previously described rafts or dendrites (Gorski et al., 1993; Sato et al., 2021). While the connection between the carbon source and flavobacterial colony expansion is not clear, the fact that we did not observe rapid expansion or raft-like structures in flavobacteria grown on PEG8000, indicates that the phenomenon is specifically attributed to the chemical attributes of pectin and not to physical properties (viscosity) of the growth medium.

While previous studies indicate that *F. johnsoniae* cannot utilize non-depolymerized cellulose or M.cellulose a sole carbon source (McBride et al., 2009), we demonstrate that colonies of *F. johnsoniae* did expand on rich (PY2) media amended with M.cellulose in microplate experiments using fluorescent microscopy. It appears that the bacteria propagated on the microcrystal surfaces without creating a dense colony structure, suggesting that the bacterium glides along the fibers without actually metabolizing them. In contrast to pectin, growth on glucose inhibited propagation of flavobacteria as previously reported (Wolkin and Pate, 1984; Imamura et al., 2018). Specifically, glucose was shown to inhibit colony spreading via MfsA which encodes a major facilitator superfamily (MFS) transporter (Imamura et al., 2018), resulting in absence of raft microstructures that are formed when grown on other carbon sources. Nonetheless, glucose did not inhibit gliding motility in general (Wolkin and Pate, 1984; Gorski et al., 1993) and spreading on glucose was recently found to be in inverse correlation to agar concentration and characterized by unique windmill-like structures under media of specific glucose-agar levels (Sato et al., 2021)

### Pectin induces colony expansion and colonization of soil and root bacteria

Pectin, but not pectin monomers (D-galacturonic acid and L-rhamnose) significantly facilitated colony expansion in a dose dependent manner. Rhizosphere and phyllosphere associated flavobacterial genomes are genetically compatible for metabolizing plant-associated polysaccharides. Specifically, their genome contains many genes encoding glycohydrolases, polysaccharide lyases, and esterases and can efficiently degrade complex biopolymers (McBride et al., 2009) and plant-associated flavobacterial genomes have an over-representation of genes involved in the metabolism of pectin (Kolton et al., 2013). Beside the ability to degrade pectin and consume it, the reduced ability of flavobacteria colonies to spread on pectin monomers (rhamnose or/and galacturonic acid) suggests that the original spatial organization of plant cell wall components are important for recognition and expansion on it. Due to the importance of pectin and other glucans in plant cell walls, we hypothesize that soil and root-associated bacteria have devised different strategies to colonize it and use it as a que for colonization and expansion on root surfaces.

For example, *Flexibacter* sp. FS-1 was not able to glide on agarose alone but did glide on agarose amended with 1% pectin (Arlauskas and Burchard, 1982). Purified Arabidopsis polysaccharides (arabinogalactan, pectin, or xylan) triggered biofilm formation in *B. subtilis* when added to rich media and induced root colonization (Beauregard et al., 2013). Similarly, addition of pectin and sucrose to the media of the PGPR *Bacillus amyloliquefaciens* strain SQY 162, increased bacterial abundance, induced biofilm formation and improved the ability of the amended bacterium to suppress tobacco bacterial wilt disease (Wu et al., 2015). In the symbiotic nitrogen fixing bacterium *Rhizobium leguminosarum*, glucomannan-mediated attachment was important for legume infection and nodulation (Williams et al., 2008).

### Growth on Pectin is bi phasic

Live-imaging fluorescent microscopy revealed that *F. johnsoniae* growth on PY2 media coated with different mono- and polysaccharides was bi-phasic in nature, with an initial phase of rapid expansion, followed by biomass production within the colonized area. This bi-phasic growth pattern, which was most pronounced on pectin, resembles previously described models in motile *E. coli*, which depicted an initial expansion phase, where “pioneer” bacteria with high velocity advance in front of the colony, followed by a second phase, where “settler” bacteria grow and replicate locally(Cremer et al., 2019; Liu et al., 2019). We hypothesize that plant-derived polysaccharides and especially pectin may serve as signal that facilitates the expansion of “pioneer” cells, and later as carbon sources that support growth of “settlers”.

### Pectin does not induce gliding-associated proteins

Despite the substantial evidence that pectin stimulated flavobacterial colony expansion, it did not induce expression of known gliding motility proteins (although not all were identified in the proteomic analyses) or proteins associated with chemotaxis, nor did it induce expression of remA, encoding the lectin binding, flavobacterial cell surface adhesin involved in gliding (Shrivastava et al., 2012). While the specific correlation between expansion on pectin and the unique flavobacterial gliding motility mechanism was not determined in this study, it is evident that the latter is required, because gliding deficient *(ΔgldJ)* flavobacterial mutants did not expand on pectin. The gliding machinery might be induced in the earlier phase of the response to pectin, in which we observed intensive bacterial motility. Alternatively, pectin might facilitate gliding in a protein expression-independent manner or induce other components of the gliding machinery not identified in our proteomic analyses. A recent study identified two flavobacterial lipoproteins linked to both biofilm formation and gliding motility (Sato et al 2021), supporting the notion that other currently hypothetical proteins may also be linked to colony spreading.

Based on proteomic and subsequent qPCR (quantitative real-time PCR) gene expression validation, we believe that the pectin-induced flavobacterial expansion observed in this study is at least partially mediated by the induction of specific TonB-associated PULs. Extrapolation of these lab-based results to flavobacterial-root interactions, suggesting a link between niche recognition, colony expansion and metabolic fitness. Similar induction of TonB and PUL was observed in marine flavobacteria as response to phytoplankton blooms characterized in decomposition of alga-derived organic matter (Teeling et al., 2012).

A few studies have previously linked Ton B proteins with motility, attachment or plant-bacterial interactions. Ton B was associated with twitching motility in *P. aeruginosa* (Huang et al., 2004), and was also found to play a role in the ability of *A. baumannii* to bind to the high-molecular weight glycoprotein fibronectin, indicating the capacity to bind to extracellular host proteins (Zimbler et al., 2013). In *Xanthomonas campestris pv. Campestris*, pectate sensed by specific TonB-dependent receptors triggered secretion of extracellular polygalacturonate. This resulted in pectin degradation and generation of oligogalacturonides (OGA) that are recognized as damage-associated molecular patterns (DAMPs), facilitating the initiation of the plant defense mechanisms (Vorhölter et al., 2012). Previous experiments showed that *Pseudomonas putida* mutants that lacked TonB, were deficient in their capacity to uptake iron and displayed impaired seed colonization, linking *TonB* to metabolic and functional fitness in plant-associated bacteria (Molina et al., 2005). The high energetic cost and substrate specificity of TonB transducers explains why the genes encoding them were induced by pectin and not constitutively expressed (Postle and Kadner, 2003; Postle, 2007). These proteins may play a pivotal role in flavobacterial-plant interactions, however, knock-out of specific *tonB* genes in *F. johnsoniae* will be challenging due to the multitude of predicted tonB genes in its genome, which suggests a high level of functional redundancy (McBride et al., 2009).

### Additional pectin-induced proteins

Pectin significantly induced several proteins in addition to TonB related proteins. These included an auxin-regulated protein, which is especially interesting since auxin is a major phytohormone responsible for plant growth and development, demonstrating again a possible connection between pectin sensing and flavobacterial-plant interactions. Interestingly, pectin resulted in substantial induction of a putative transcriptional regulator, suggesting pectin-induced regulation of additional genes. Knocking out or silencing this regulator can shed light on this pectin-induced downstream response, and its potential role in flavobacterial-plant interactions.

The fact that gene expression (qPCR) did not completely correlate with the proteomic data might be explained by post-translational modifications or regulation affecting protein stability, degradation and complex formation, as shown in similar cases (Flory et al., 2006; Liu et al., 2016). Alternatively, this discrepancy may be explained by a recently proposed model showing that in growing cells, mRNA can saturate ribosomes thereby limiting translation, resulting in an increase in the protein-to-DNA ratio (Lin and Amir, 2018).

### Summary

To summarize, we found that pectin, a prominent plant cell wall polysaccharide, facilitates expansion of flavobacteria on solid surfaces, even in the presence of nutrient-rich media. We postulate that pectin may enhance the capacity of flavobacteria to efficiently colonize and proliferate on plant surfaces. The interaction between pectin (and potentially other root glycans) and flavobacteria is mediated by induction of TonB/SusC operons and other associated PULs that facilitate metabolism of pectin. Thus, in the root environment, plant cell wall polysaccharides, and specifically pectin, may not only serve as a nutrient source for flavobacteria, but also as a potential environmental cue for colonization and rapid expansion along the root surface.

## Acknowledgments

We thank Prof. M. J. McBride from the University of Wisconsin–Milwaukee for generously providing plasmids and *F. johnsoniae* strains. We would like to thank Alla Usyskin-Tonne, for her help with the proteomics functional annotation and Eduard Belausov for his help with the binocular based imaging. This paper was published at bioRxiv as preprint, doi: https://doi.org/10.1101/2020.06.26.174714.

