## Supplementary data for "Pectin induced colony expansion of soil-derived Flavobacterium strains"

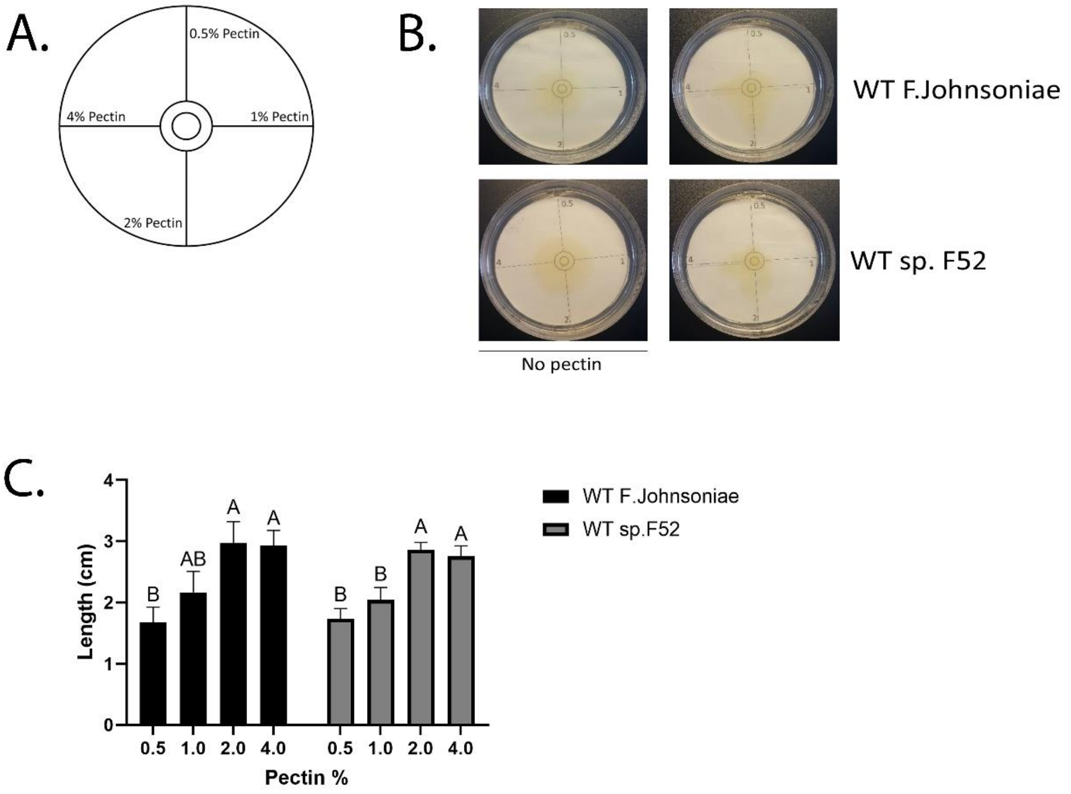


**FigS1– Effect of pectin concentration on Flavobacteria colony expansion.** (A) Schematic representation of PY2 agar plates streaked with 0.5, 1, 2, 4% pectin as indicated. (B) Wild type (WT) or gliding mutant (ΔgldJ) of *F. johnsoniae*, sp. *F52*, flavobacteria strains were inoculated in the center of PY2 plates streaked with lines of various pectin concentrations (as indicated in A.) or PY2 agar plates streaked with DDW as a control. Bacteria were grown for 48hr at $30^{O}$C and the distance of the bacterial colony from the center outwards was measured using Fiji. $p<0.05$ by Tukey HSD. N=4. (C) Distance of WT *F. johnsoniae*, *Flavobacterium* sp. F52 strains from the plate center outwards measured on PY2 agar covered with the indicated concentration of pectin. Presented bars show mean and SE values. $p<0.05$ by Tukey HSD.

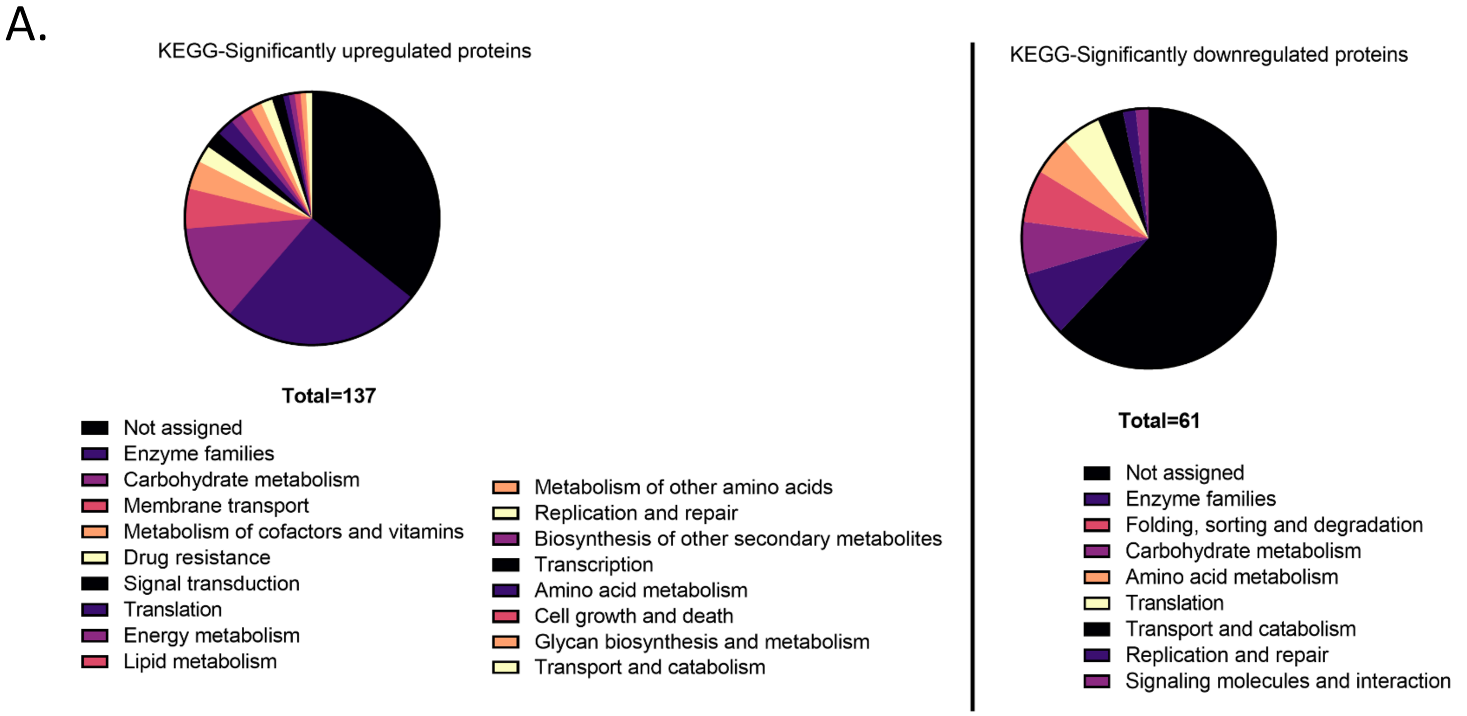
A.
KEGG-Significantly upregulated proteins
Total=137
Not assigned
Enzyme families
Carbohydrate metabolism
Membrane transport
Metabolism of cofactors and vitamins
Drug resistance
Signal transduction
Translation
Energy metabolism
Lipid metabolism
Metabolism of other amino acids
Replication and repair
Biosynthesis of other secondary metabolites
Transcription
Amino acid metabolism
Cell growth and death
Glycan biosynthesis and metabolism
Transport and catabolism
KEGG-Significantly downregulated proteins
Total=61
Not assigned
Enzyme families
Folding, sorting and degradation
Carbohydrate metabolism
Amino acid metabolism
Translation
Transport and catabolism
Replication and repair
Signaling molecules and interaction

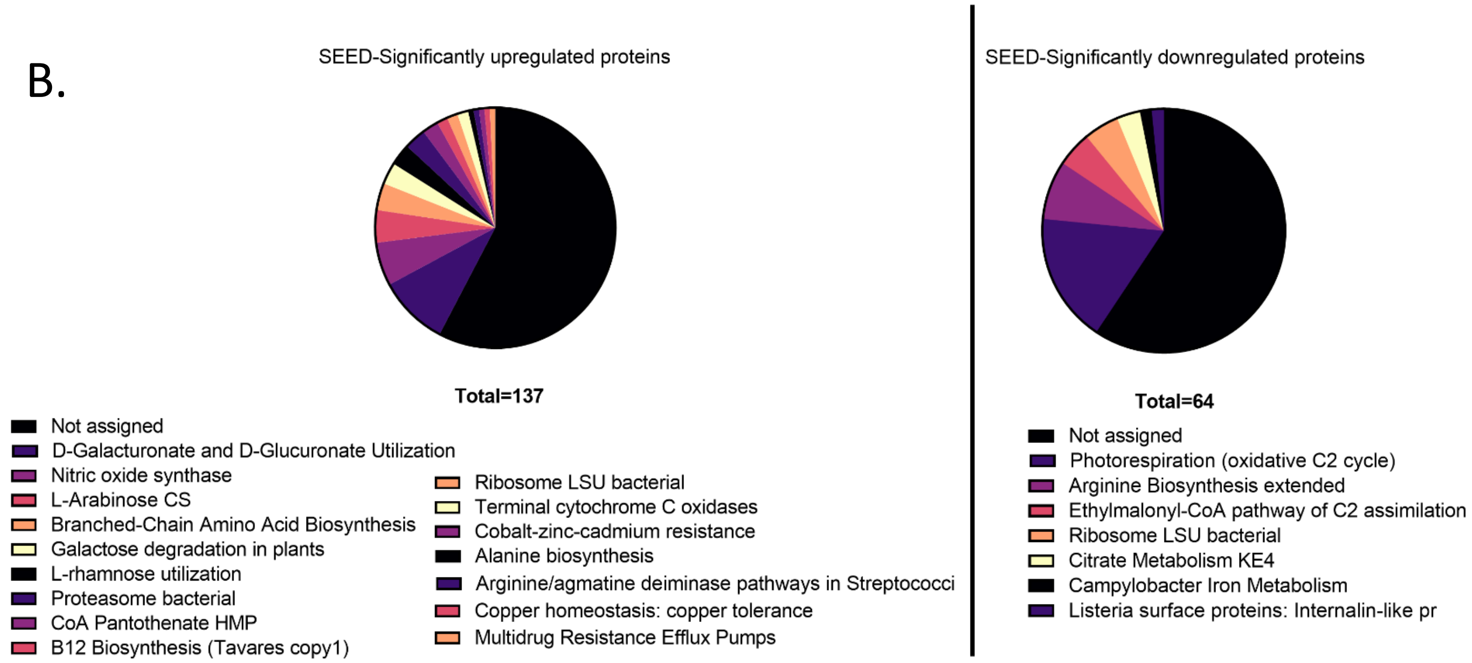
B.
SEED-Significantly upregulated proteins
Total=137
Not assigned
D-Galacturonate and D-Glucuronate Utilization
Nitric oxide synthase
L-Arabinose CS
Branched-Chain Amino Acid Biosynthesis
Galactose degradation in plants
L-rhamnose utilization
Proteasome bacterial
CoA Pantothenate HMP
B12 Biosynthesis (Tavares copy1)
Ribosome LSU bacterial
Terminal cytochrome C oxidases
Cobalt-zinc-cadmium resistance
Alanine biosynthesis
Arginine/agmatine deiminase pathways in Streptococci
Copper homeostasis: copper tolerance
Multidrug Resistance Efflux Pumps
SEED-Significantly downregulated proteins
Total=64
Not assigned
Photorespiration (oxidative C2 cycle)
Arginine Biosynthesis extended
Ethylmalonyl-CoA pathway of C2 assimilation
Ribosome LSU bacterial
Citrate Metabolism KE4
Campylobacter Iron Metabolism
Listeria surface proteins: Internalin-like pr

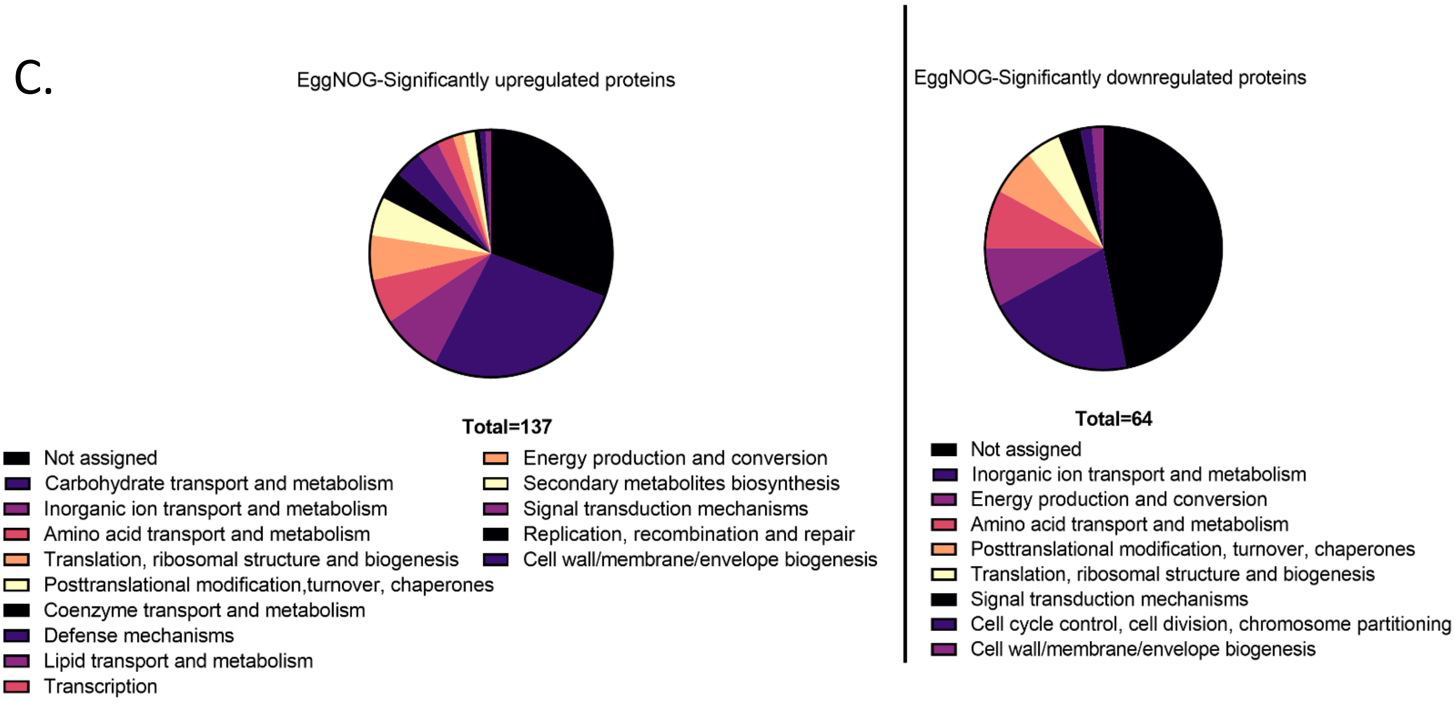


**FigS2– Annotation of proteins upregulated and down regulated in response to pectin in *F. johnsoniae***

Proteins that were significantly altered and had fold change higher then 2 (left panel) or lower then 0.5 (right panel) between flavobacteria grown on PY2 with pectin or DDW were annotated using either: (A) KEGG (B) Seed (C) EggNog.

### Supplementary Materials and Methods

#### Functional annotation of significantly induced and repressed proteins

Proteins showing at least two-fold increase or decrease in PY2 media amended with pectin relative to the PY2 media amended with DDW (control) were screened against the non-redundant (nr) NCBI protein database using DIAMOND v0.9.24.125 (Buchfink et al., 2015) to assign functional annotations. Results were then uploaded to MEGAN Ultimate edition software v6.15.2 (Buchfink et al., 2014). For functional annotation, the Kyoto Encyclopedia of Genes and Genomes (KEGG), SEED (Overbeek et al., 2014), and EggNOG (Based on Clusters of Orthologous groups – COG extension) databases were used.

#### Operon mapping of TonB-associated gene cluster

A TonB gene cluster that was significantly upregulated in the pectin samples (peg4455-4469 by RAST, WP_044048002.1-OXE95634.1 by NCBI annotation, bases 5125269-5130351 in *F. johnsoniae* UW101, complete genome Sequence ID: CP000685.1) was screened for the presence of operons using two independent tools: Operon mapper (Taboada et al., 2018), and FGENESB Suite of Bacterial Operon-Softberry Inc. (Tyson et al., 2004).

**Table S1A- Significantly upregulated proteins of *F. johnsoniae* grown on PY2 covered with pectin vs. DDW**

| NCBI annotaion | gene_number (RAST annoation) | Pectin/DDW: P-Value | Pectin/DDW: Fold change | DNA seq | RAST functional annotation |
|---|---|---|---|---|---|
| WP_012024292.1 | 2323 | 0.00669 | **585.7** | atgaaaaaatcaattctaattctattcattttaaccagctttt | hypothetical protein |
| WP_012022292.1 | 192 | 0.00148 | **203.9** | atgaaagcaaaatatatatttagaggactaatagccgta | hypothetical protein |
| WP_012023625.1 | 1618 | 0.01686 | **128.9** | atgttgggattaaaattagctacagaccctcgctgggtta | tRNA-(ms[2]io[6]A)-hydroxylase (EC 1.-.-.-) |
| WP_012024113.1 | 2127 | 0.04239 | **84.3** | atgttacaactaaaaatcaaaaaaacagccattattttg | Beta-glucosidase (EC 3.2.1.21) |
| WP_012025397.1 | 3584 | 0.00002 | **38.6** | atgaagaatgtaactatcaaattgttagcagctcttgtatt | Alkaline phosphatase (EC 3.1.3.1) |
| WP_012023204.1 | 1170 | 0.00388 | **29.9** | atgaaaaattatgtcataggattagactacggaacaga | Ribulokinase (EC 2.7.1.16) |
| WP_012023111.1 | 1074 | 0.00438 | **26.7** | atgtcaattaaatcgatagcagcaaagatatttgcccgc | putative auxin-regulated protein |
| WP_012023206.1 | 1172 | 0.00564 | **25.8** | atgatagatatatctcaaaaagaagtatggtttgtagtag | L-arabinose isomerase (EC 5.3.1.4) |
| WP_012024544.1 | 2583 | 0.00180 | **23.1** | atgaaactaaattttctaaaatcaattgcactgcttggtat | SclB protein |
| WP_012022294.1 | 194 | 0.00139 | **22.1** | atgaattttaaagatttattaaacaaaggagcaaatcgtt | TonB family protein / TonB-dependent receptor |
| WP_012022711.1 | 646 | 0.04622 | **19.3** | atgaaaacaataaataaacttatacttattcttctttctcta | hypothetical protein |
| WP_012022637.1 | 570 | 0.00997 | **14.4** | atgaaaaaatcaattttaaatttcgttacagttgttgctttat | FIG00649626: hypothetical protein |
| WP_012025446.1 | 3634 | 0.02934 | **12.9** | atgattttttctttaattataccagtgtataatcgtccagatga | Glycosyltransferase |
| WP_012023596.1 | 1589 | 0.00918 | **12.8** | atgaaaaaaatgctacttattcttgcattgtcagtttttggct | FIG00657157: hypothetical protein |
| WP_012022876.1 | 819 | 0.00389 | **12.1** | atgctagacgaaaccccaaaacgatttgatagaattgtt | transcriptional regulator, putative |
| SHH12854.1 | 4459 | 0.00449 | **11.9** | atgttaggaatgattgttacaagcttaagttcttgcagcaa | RagB/SusD domain protein |
| WP_012023313.1 | 1287 | 0.00123 | **11.4** | atgaaaaaaaacctactttttctgactatgcttctaattgct | Endo-1,4-beta-xylanase A precursor (EC 3.2.1.8) |
| WP_012026225.1 | 4456 | 0.01930 | **11.2** | atgataaagaaaagccattttagttttgcccaaatcatgc | Rhamnogalacturonides degradation protein RhiN |
| WP_012022957.1 | 905 | 0.01684 | **10.1** | atgagaacgtgtcttgaatgctctgaaaacttgtaggca | FIG00930353: hypothetical protein |
| WP_012026229.1 | 4460 | 0.00035 | **8.4** | atgaacattcaaaaatcattaaagaaaaaaataaaata | TonB family protein / TonB-dependent receptor |
| WP_012022293.1 | 193 | 0.00954 | **8.4** | atgaaacataaattaataatagcaggattgataatttcaa | putative outer membrane protein, probably involved in nutrient binding |
| WP_012026233.1 | 4464 | 0.00003 | **7.7** | atggaaaaaattaacagatcaaattcagagttttcaaac | Altronate oxidoreductase (EC 1.1.1.58) |
| WP_012026234.1 | 4465 | 0.00005 | **6.8** | atggcaaaatattcaagaatagaggttgcgcagacaa | 4-hydroxy-2-oxoglutarate aldolase (EC 4.1.3.16) @ 2-dehydro-3-deoxyphosphogluconate aldolase (EC 4.1.2.14) |
| WP_012026236.1 | 4467 | 0.00006 | **6.4** | atgagcgcaaatacattcatacacgacaatttttattaga | Uronate isomerase (EC 5.3.1.12) |
| WP_012022307.1 | 209 | 0.03901 | **6.1** | atgtcagatacaatcgaaaaaattaaatgccttattatag | Thioredoxin reductase (EC 1.8.1.9) |
| WP_012026223.1 | 4454 | 0.00058 | **6.0** | atgaaaaatataaaaatcgttattcttctcctttccgtttttc | Beta-xylosidase (EC 3.2.1.37) |
| WP_011921584.1 | 72 | 0.03454 | **5.9** | atggtaaaatttggatatacaattttgtacgttgaagacgt | related lyases |
| WP_012026232.1 | 4463 | 0.00037 | **5.7** | atggcagcgcagaaaaaattgataaaagttcacccaa | Altronate hydrolase (EC 4.2.1.7) |
| SHL59770.1 | 2398 | 0.02051 | **5.5** | atgtcacaaaataatatcattttaacaacaggaacatat | Regulator of nucleoside diphosphate kinase |
| WP_012026691.1 | 4957 | 0.01793 | **5.5** | atggaaaaatggaaacaaaatttaatcgccgcaattct | TonB-dependent receptor, putative |
| WP_089477413.1 | 4508 | 0.04171 | **5.2** | atgaagaaaatagcatgtattgcgcttttagctttactattt | hypothetical protein |
| OXE95634.1 | 4469 | 0.00400 | **4.8** | atgacaaaatatagttcaagatacgcgtcaagtccaga | 4-deoxy-L-threo-5-hexosulose-uronate ketol-isomerase (EC 5.3.1.17) |
| WP_012026094.1 | 4323 | 0.04956 | **4.7** | atgaaattaatacgttttggagaaatcggaaaagaaaa | 5-oxopent-3-ene-1,2,5-tricarboxylate decarboxylase (EC 4.1.1.68) |
| WP_012026239.1 | 4470 | 0.00163 | **4.6** | atgagtcaaaccgatgctgttacagtaaatcaaaccgtt | Hexuronate transporter |
| WP_044048002.1 | 4455 | 0.00000 | **4.6** | atgtcggttttattagtttgcgtaatggctttaacaggctgta | Rhamnogalacturonides degradation protein RhiN |
| WP_012026071.1 | 4300 | 0.00094 | **4.4** | atgaaaaagaacatcatcatatcctctttatttctttcaacc | Putative glycosyl hydrolase of unknown function (DUF1680) |
| WP_012023778.1 | 1772 | 0.00746 | **4.3** | atgaactacggtaaagaatttaaaaattttgctacaaag | ATP-dependent Clp protease proteolytic subunit (EC 3.4.21.92) |
| WP_044047683.1 | 2297 | 0.01650 | **4.1** | atgagagaaaatgacatgttagaaaaagaagagattt | FIG00655294: hypothetical protein |
| WP_012023203.1 | 1169 | 0.00112 | **4.0** | atgaaaaaaacactactactattgtttgttacaatttgtata | Alpha-N-arabinofuranosidase 2 (EC 3.2.1.55) |
| WP_012023735.1 | 1729 | 0.00019 | **3.8** | atggaaatgacattaacaacaagtgatgagcaggtaa | Alternative cytochrome c oxidase polypeptide CoxO (EC 1.9.3.1); Cytochrome c oxidase, subunit III (EC 1.9.3.1) |

| | | | | | |
|---|---|---|---|---|---|
| WP_091490394.1 | 1180 | 0.00210 | **3.8** | atggcttttgaattaccacaattaccttatgcatacgatgc | Manganese superoxide dismutase (EC 1.15.1.1) |
| WP_044048511.1 | 4466 | 0.00013 | **3.8** | atgagtagagtagttgcatttggagaaatcatgctgcgtt | 2-dehydro-3-deoxygluconate kinase (EC 2.7.1.45) |
| WP_012026231.1 | 4462 | 0.00004 | **3.7** | atgagcgaaaaagtaaccatttatgatattgccgaaaa | Hexuronate utilization operon transcriptional repressor ExuR |
| WP_012026237.1 | 4468 | 0.01289 | **3.5** | atgacaaacttatttgacataaaaggaaaagttgccctt | 2-deoxy-D-gluconate 3-dehydrogenase (EC 1.1.1.125) |
| WP_012024354.1 | 2387 | 0.02187 | **3.4** | atgttccagaaaattactcttttaacgcttgtattagctgctg | hypothetical protein |
| WP_012026430.1 | 4683 | 0.04473 | **3.3** | atgacaatgtattcattcgatataaaaaataaaaaccgg | Probable Co/Zn/Cd efflux system membrane fusion protein |
| WP_012024436.1 | 2473 | 0.00884 | **3.2** | atgaaaatacaagatattcaggcaattcaattattgcttg | oligoribonuclease A, Bacillus type |
| WP_052295191.1 | 3036 | 0.04623 | **3.2** | atgaaaaatataaaaataacactttcattattattggtgat | Putative outer membrane protein, probably involved in nutrient binding |
| WP_044047803.1 | 3435 | 0.01612 | **3.1** | atgtcaagcaaagaagcatataaaggtgatcttgatac | FIG01101450: hypothetical protein |
| WP_012026188.1 | 4419 | 0.00004 | **3.1** | atgataatcggatcaaaccatattgaatctcataacgag | Predicted L-rhamnose isomerase RhaI (EC 5.3.1.14) |
| WP_012025223.1 | 3406 | 0.00852 | **3.0** | atgagcattaaaaacaattcatttttcatctctattttatcac | Probable RND efflux membrane fusion protein |
| WP_012026482.1 | 4737 | 0.00920 | **2.9** | atgagaatcatttcaggaaaatacaaagggcgccgga | 16S rRNA (guanine(966)-N(2))-methyltransferase (EC 2.1.1.171) |
| WP_012026068.1 | 4297 | 0.02085 | **2.9** | atgaaaacaaaatttaataaatatatatcactggcacttt | putative outer membrane protein, probably involved in nutrient binding |
| WP_012026259.1 | 4490 | 0.00199 | **2.8** | atgaatttaatacgttttgcactccgcaaacccatttccatt | RND multidrug efflux transporter; Acriflavin resistance protein |
| WP_012025845.1 | 4060 | 0.01054 | **2.7** | atgacagtacatcataatattaccaatggatacctgagta | Endo-1,4-beta-xylanase A precursor (EC 3.2.1.8) |
| WP_012026180.1 | 4411 | 0.04757 | **2.6** | atgaaaaaatttagttattttattagcagttgcaattcagttaa | Copper homeostasis protein CutF precursor / Lipoprotein NlpE involeved in surface adhesion |
| SEP16691.1 | 568 | 0.04429 | **2.6** | ttgctatttatgaaaaaaacacaaactaagattttattagt | Response regulator receiver:Transcriptional regulatory protein, C- terminal |
| SHF95927.1 | 1833 | 0.02412 | **2.5** | ttgccaactgaaaatctgaaaaatatgatcatccaaaa | Methionine aminopeptidase (EC 3.4.11.18) |
| WP_011921605.1 | 94 | 0.00879 | **2.5** | atgagccatttagacgatatattaaaatcacgccgcgat | Nicotinate-nucleotide--dimethylbenzimidazole phosphoribosyltransferase (EC 2.4.2.21) |
| WP_012025858.1 | 4074 | 0.00397 | **2.5** | atgaaacgatatatttttagaaatttaataataggatctgta | RagB/SusD domain protein |
| WP_012023590.1 | 1583 | 0.00139 | **2.4** | atgaaacaagaagaactacaagccatagcctctcaat | Biotin synthesis protein BioC |
| WP_012022472.1 | 385 | 0.04716 | **2.4** | atgttgcaattaaatgtaaagaacgaaacatctaggctt | NG,NG-dimethylarginine dimethylaminohydrolase 1 (EC 3.5.3.18) |
| WP_012026165.1 | 4396 | 0.00089 | **2.3** | atggaacaaacattaagatggttcggaccaaatgatcc | Mannonate dehydratase (EC 4.2.1.8) |
| WP_012026655.1 | 4918 | 0.00377 | **2.3** | atgaatacagcagattttactacaacaatacttgtgagcc | hypothetical protein |
| WP_012024856.1 | 2932 | 0.00615 | **2.3** | atgaataccaaaaatagtaccatcagtctagaaacttat | Cysteine desulfurase (EC 2.8.1.7) |
| WP_012023797.1 | 1792 | 0.03633 | **2.2** | atgagaaaacttgagaatagcgaattagaaagaaaat | TRNA/rRNA methyltransferase |
| WP_012022623.1 | 556 | 0.01067 | **2.2** | atgccattttcagaattatttgataacgaattcaaacaaa | FIG00651957: hypothetical protein |
| WP_012026675.1 | 4939 | 0.02216 | **2.2** | atgaaaaaaatctacttcttaattgcattattctgcaccgtt | hypothetical protein |
| WP_012023198.1 | 1163 | 0.03806 | **2.2** | atgaaaaattttgacattaacgaagatccacacagacg | Galactose-1-phosphate uridylyltransferase (EC 2.7.7.10) |
| WP_012025504.1 | 3697 | 0.03226 | **2.1** | atgattagaatttcagttttatttttagcttttgcttttttagttgt | hypothetical protein |
| WP_012022715.1 | 651 | 0.00014 | **2.1** | atgaaaataaaactatacgccctttctacagttcttacgc | hypothetical protein |
| WP_012025247.1 | 3431 | 0.01325 | **2.1** | atgtcgtttttaacatcattaacagtagcaggaaaagatt | hypothetical protein |
| WP_012024304.1 | 2335 | 0.00569 | **2.1** | atgatttcatttctggatctaaaaaaaataaatcagccata | Aminotransferase |
| WP_012026057.1 | 4285 | 0.02152 | **2.1** | atgggattgaaaattaaaataactgcgatttgttttggaat | putative large secreted protein |
| WP_012024919.1 | 3003 | 0.00072 | **2.1** | atggcaaattatttcaacacactaccacttagattacaat | Ketol-acid reductoisomerase (EC 1.1.1.86) |
| WP_012025852.1 | 4068 | 0.00074 | **2.1** | atgaaagtactctataacctattagcgattttttaattcttac | glycoside hydrolase, family 43 |
| WP_044047537.1 | 1189 | 0.02710 | **2.0** | atggcattaaggcgagttgttgtaacaggattaggtgca | 3-oxoacyl-[acyl-carrier-protein] synthase, KASII (EC 2.3.1.41) |
| WP_012026676.1 | 4940 | 0.02970 | **2.0** | atgtcaacagcaaaaaaagattataaaagaatcacaa | 3-methyl-2-oxobutanoate hydroxymethyltransferase (EC 2.1.2.11) |
| WP_012022972.1 | 922 | 0.03873 | **2.0** | atgaaaaaagttatcatcacattattgtttttagtatcatttg | hypothetical protein |
| WP_012026682.1 | 4946 | 0.03967 | **2.0** | atgaaattagacatattagcctttggcgcacatccggatg | LMBE-RELATED PROTEIN |

| | | | | | |
|---|---|---|---|---|---|
| WP_012026316.1 | 4551 | 0.04159 | **2.0** | atgaaaaaaatagttgaataccgcaagttactaaacgt | chaperone with DnaK; heat shock protein |
| WP_007803649.1 | 405 | 0.02033 | **2.0** | atggtacaacaagaatcaagactaaaagtagcagata | LSU ribosomal protein L14p (L23e) |

**Table S1B- Significantly downregulated proteins of *F. johnsoniae* grwon on PY2 covered with pectin vs. DDW**

| NCBI annotation | gene_number (RAST annoation) | Pectin/DDW: P-Value | Pectin/DDW: Fold change | DNA seq | RAST functional annotation |
|---|---|---|---|---|---|
| WP_011921578.1 | 2219 | 0.01329 | **0.0** | atggattcagaattaaattcgaaagtcaatctgcaggaa | Possible Galanin |
| WP_012022355.1 | 1796 | 0.01684 | **0.0** | atgaaaaaagtatttttaagtttagctgttgttgctgttttaac | hypothetical protein |
| WP_012022481.1 | 2768 | 0.00263 | **0.1** | atgaaaaatacagaaacaaaaaatactaaagatacg | protein of unknown function DUF892 |
| WP_012022486.1 | 2790 | 0.02104 | **0.1** | atgaaaacagatacaaagggaaaccagcgtcagga | hypothetical protein |
| WP_073099567.1 | 2317 | 0.00190 | **0.1** | atgaaaaagttatattatacagcaaacgtactacttattct | ErfK/YbiS/YcfS/YnhG family protein |
| WP_012022550.1 | 3180 | 0.01144 | **0.2** | atggagccacttatttattatccgacttttgagccgccaac | hypothetical protein |
| WP_012023095.1 | 3303 | 0.00112 | **0.2** | atgaaaaattttcaggatgaaaagcagcgagatctgct | Catalase (EC 1.11.1.6) |
| WP_012023270.1 | 4631 | 0.01186 | **0.2** | atgaatcttattaaaagaaacggaagccatgtgccggc | Small heat shock protein |
| WP_012022355.1 | 263 | 0.00189 | **0.2** | atgaaaacaaaattcttaaaactttcgcttttagcactatt | hypothetical protein |
| WP_044048264.1 | 1817 | 0.00385 | **0.2** | atgatacatacagatgaaaccacacaagaagctgtta | hypothetical protein |
| WP_012023815.1 | 2422 | 0.01124 | **0.2** | atgactccaaatatcggaatatctcccgccaacttaaaa | Non-specific DNA-binding protein Dps / Iron-binding ferritin-like antioxidant protein / Ferroxidase (EC 1.16.3.1) |
| WP_012023822.1 | 3937 | 0.00564 | **0.2** | atgaatcttatcaaaagaaatgcaaaccagcatcgtgc | Small heat shock protein |
| WP_012023992.1 | 1243 | 0.02321 | **0.2** | ttgacacagttaaatctgtctaaaaacagagctttaacct | Internalin-like protein (LPXTG motif) Lmo0331 homolog |
| WP_012024192.1 | 3531 | 0.00304 | **0.2** | atgggagatcataaagaccttactgaagaatttgctgtc | Pyridoxamine 5'-phosphate oxidase (EC 1.4.3.5) |
| WP_012024286.1 | 5085 | 0.00000 | **0.3** | atggaatacaaagtagtaccattcgtagcttcaatcgat | hypothetical protein |
| N/A | 5088 | 0.00065 | **0.3** | ttgggaggaattgtttttgcaactggagctttggctgggat | hypothetical protein |
| WP_012024386.1 | 3238 | 0.02224 | **0.3** | atggaacacgcaacacagcagtattcaaaagaggta | protein of unknown function DUF305 |
| WP_012024486.1 | 3304 | 0.02676 | **0.3** | atgaaaaattcagaaaaaccaaataatacaaaagctg | protein of unknown function DUF892 |
| WP_012024493.1 | 5084 | 0.04047 | **0.3** | atgtttaaaaaagtaataattgctgaagatatcgacgca | hypothetical protein |
| WP_012024711.1 | 4786 | 0.02762 | **0.3** | atgaaagtaatattgattgttttgtgttttttacggctggtttt | hypothetical protein |
| WP_012024727.1 | 1809 | 0.00592 | **0.3** | atgaaaaaccaattagaaattaccgaaacggcaatgg | Malate synthase (EC 2.3.3.9) |
| WP_012025066.1 | 4223 | 0.00011 | **0.3** | atgaactataacaaacttttactttgtcttttatttacaatgca | hypothetical protein |
| SHK65708.1 | 2530 | 0.00116 | **0.3** | atgaaaaataaattactttggtttttagctgttttcactgtttc | Mechanosensitive ion channel family protein |
| WP_012025127.1 | 3541 | 0.00945 | **0.3** | atgaaaaagatactattagccagtaaagcaattttagga | putative exported protein |
| WP_012025128.1 | 1810 | 0.01642 | **0.3** | atgaaaacaacagaagacagaattcaggaattgatta | Isocitrate lyase (EC 4.1.3.1) |
| WP_012025137.1 | 5248 | 0.00410 | **0.3** | atgagaaatataaaaatagctgcggcagttacattatta | hypothetical protein |
| WP_012025282.1 | 407 | 0.01798 | **0.3** | atgaaacaatcagaaataaaagatctttctgcagcgga | LSU ribosomal protein L29p (L35e) |
| WP_012025337.1 | 476 | 0.01318 | **0.3** | atgagtttcaccatttcttcagataaaaagaaatcattact | Ferric siderophore transport system, periplasmic binding protein TonB |
| WP_012025345.1 | 66 | 0.03714 | **0.3** | atgaaattatacaaaatagcaaccgttcaatatgtaaat | Cell division inhibitor |
| WP_073409277.1 | 437 | 0.03784 | **0.4** | atgtctgaaattgcccttatttcagctagaaaaaaccgac | Magnesium and cobalt efflux protein CorC |
| WP_012025413.1 | 3314 | 0.01428 | **0.4** | atggcaacaacttacaaaacgcctggagtatatgttga | Phage tail sheath protein FI |
| WP_012025418.1 | 3606 | 0.01249 | **0.4** | atgaaaaaagtagtattagcttatagcggaggattagat | Argininosuccinate synthase (EC 6.3.4.5) |
| WP_012025595.1 | 2414 | 0.01205 | **0.4** | atgaaagcatctagcacaattttaggttttgcagcagcg | Gas vesicle protein |
| WP_012025730.1 | 4990 | 0.00398 | **0.4** | atgcataattatttagaaacaaatttaaaaattgaaacta | hypothetical protein |
| WP_012026000.1 | 1058 | 0.01935 | **0.4** | atggcatttgaagagttaagagaaaatactgaaaaaat | FIG00654500: hypothetical protein |
| WP_012026385.1 | 3793 | 0.00828 | **0.4** | atgagtttaggtgatttatttccagacaatctgaaagaag | hypothetical protein |
| WP_012026531.1 | 4860 | 0.01204 | **0.5** | atggaaaacaaacccgaagaaattactcctgagaaa | Low-complexity acidic protein, XCC2875 type |
| WP_089477404.1 | 3601 | 0.02595 | **0.5** | atgaactacatctcaataaaagatatcgactcattatca | Ornithine carbamoyltransferase (EC 2.1.3.3) |
| ABQ07757.1 | 2523 | 0.00426 | **0.5** | atgaaaaaagtactaacactaacggcccttgtttttacg | hypothetical protein |
| WP_012026811.1 | 1992 | 0.00004 | **0.5** | atgaaaaaattatttatcgcagcaatgttatttgttggaata | hypothetical protein |
| WP_044048134.1 | 3524 | 0.02029 | **0.5** | atgaatggtaactttaaaagaaatctgcttataagttcatt | Signal transduction histidine kinase |
| WP_012026814.1 | 3466 | 0.00608 | **0.5** | atgataaaatacagcccacaaaattactttaaatgcctg | Antifreeze protein |
| WP_012026963.1 | 397 | 0.01510 | **0.5** | atggctaaattattagtaaaacaagtaagaagcaagat | LSU ribosomal protein L30p (L7e) |

**Table S2- Fold change of proteins related to *F. johnsoniae* gliding machenary in pectin vs. DDW**

| Protein | NCBI annotation | Gene Sequence | # unique peptides | Pectin/DDW: P-Value | Pectin/DDW: Fold change | Annotation by RAST |
|---|---|---|---|---|---|---|
| RemC | WP_012022324.1 | atgaaaagaaaaatactttttcttgg | 2 | 0.08363 | **13.0** | Glycosyltransferase (EC 2.4.1.-) |
| Wzc | WP_012022466.1 | atgttagatataaaagattttccatt | 19 | 0.38789 | **0.9** | Tyrosine-protein kinase Wzc (EC 2.7.10.2) |
| Wza | WP_012022467.1 | atgacaaaaaatagcttttatatact | 5 | 0.14301 | **1.2** | Polysaccharide export outer membrane protein |
| SprF | WP_012023064.1 | atgatgttatctaaaaaaattattac | 4 | 0.06417 | **1.2** | FIG00654737: hypothetical protein |
| SprB | ABL60886.1 | atgaaaaaacctactattttaagaa | 27 | 0.00858 | **1.3** | internalin, putative |
| SprD | WP_012023066.1 | atgaagaaaattttactattcataac | 28 | 0.06564 | **0.6** | TolA protein |
| SprC | WP_012023067.1 | atgattcaaaaaactactttatctttt | 7 | 0.62616 | **0.9** | FIG00654018: hypothetical protein |
| RemG | WP_012023069.1 | atgaaaaaaattactcaaatgatct | 6 | 0.19823 | **1.1** | FIG00653638: hypothetical protein |
| RemF | WP_012023070.1 | atgaaagggaaaattattttatttag | 1 | 0.25834 | **0.7** | FIG00655764: hypothetical protein |
| GldE | WP_012023105.1 | atgaaaatttcgataggaaacgac | 7 | 0.09691 | **0.7** | Ribose 5-phosphate isomerase B (EC 5.3.1.6) |
| SprE | WP_012023137.1 | ttgtataatggaggtcttggacttga | 13 | 0.06601 | **0.8** | TPR domain protein |
| SprT | WP_012023545.1 | atgagaaaaattgtaatcgtcatttt | 4 | 0.49504 | **1.1** | PorT protein |
| GldA | WP_012023594.1 | atgtcgatagaagtaaacagtata | 4 | 0.54640 | **1.1** | ABC transporter, ATP-binding protein |
| GldD | WP_012023616.1 | atgttaaaaaaaatcctttcagtaa | 2 | 0.25781 | **1.6** | GldD |
| PorV | WP_012023631.1 | atgaaaaaaatatcgcttctattaat | 15 | 0.19036 | **1.4** | FIG00650346: hypothetical protein |
| SprA | WP_012023729.1 | atggagttagaaaatcctccaagc | 28 | 0.03250 | **1.1** | FIG00648977: hypothetical protein |
| RemB | WP_012023733.1 | gtgaataagttatttttatcttttatcat | 4 | 0.10729 | **2.1** | FIG00654784: hypothetical protein |
| GldB | WP_012023869.1 | atgaaaatgtatcgctttgtagtggt | 6 | 0.07962 | **2.7** | GldB |
| GldC | WP_012023870.1 | atgtcaaatacaataaaatcagaa | 1 | 0.83912 | **0.9** | GldC |
| GldJ | WP_012023929.1 | atgaagaagtttattgcatttgcagc | 19 | 0.05145 | **0.9** | GldJ |
| GldL | WP_012023930.1 | atggcattattaagtaaaaaagtta | 23 | 0.27486 | **1.1** | FIG00648934: hypothetical protein |
| GldM | WP_012023931.1 | atgtatctggttttcatcgcaatgtta | 53 | 0.00280 | **1.1** | FIG00649784: hypothetical protein |
| GldN | WP_012023932.1 | atgaaagtaagaaatttttaatagc | 15 | 0.96310 | **1.0** | GldN |
| GldO | WP_012023933.1 | atgaaagtaaggaatttttaattgc | 15 | 0.02686 | **0.7** | GldN |
| GldI | WP_012024435.1 | atgaactacttaaaaatcagcattta | 4 | 0.47280 | **1.0** | GldI |
| GldG | WP_012024782.1 | atgaaagcatctaataaattaaatc | 11 | 0.64396 | **1.0** | gliding motility protein GldG |
| RemI (RemG-paralog) | WP_012025180.1 | atgatttgtttactgctgacagctgcc | 7 | 0.34787 | **1.8** | FIG00653638: hypothetical protein |

**Table S3- Pectin induced TonB regulated gene clusters**

Cluster 1

| NCBI annotation | # unique peptides | Pectin/DDW: P-Value | Pectin/DDW: Fold change | Gene Sequence | Signal Peptide? | RAST annotaion |
|---|---|---|---|---|---|---|
| WP_012026221.1 | 1 | 0.05681 | **18.2** | atgaaatatttacttctattattaatttcctatgtttcatttgctcagcaaaataattttccaagagcaaa | Y | Polygalacturonase (EC 3.2.1.15) |
| WP_012026223.1 | 5 | 0.00058 | **6** | atgaaaaatataaaaatcgttattcttctcctttccgtttttcttttcaaaaaaatacggcacaagcc | Y | Beta-xylosidase (EC 3.2.1.37) |
| WP_044048002.1 | 11 | 0 | **4.6** | atgtcggttttattagtttgcgtaatggctttaacaggctgtaaagtaacttcacaagaaacggcaa | Y | Rhamnogalacturonides degradation protein RhiN |
| WP_012026225.1 | 5 | 0.0193 | **11.2** | atgataaagaaaagccattttagttttgcccaaatcatgcttataagtatgatgctgttttcattgaat | Y | Rhamnogalacturonides degradation protein RhiN |
| WP_012026226.1 | 2 | 0.08763 | **7.4** | atgaactataaaaaaacagtctttgcatttttcctgttgtacagcatttttactgtaggtcagaacaat | Y | Rhamnogalacturonides degradation protein RhiN |
| SHH12854.1 | 7 | 0.00449 | **11.9** | atgttaggaatgattgttacaagcttaagttcttgcagcaattatatagaggaggaaagtttatcaa | N | RagB/SusD domain protein |
| WP_012026229.1 | 28 | 0.00035 | **8.4** | atgaacattcaaaaatcattaaagaaaaaaataaaatacaatcttgtatttttatttttcctgaatttt | Y | TonB family protein / TonB-dependent receptor |
| WP_012026230.1 | 2 | 0.13578 | **2.8** | ttgactatatcatgctcaaaaaaaattactcctgaagctgaaactgatccgtggaaaactatgga | N | Polygalacturonase (EC 3.2.1.15) |
| WP_012026231.1 | 6 | 0.00004 | **3.7** | atgagcgaaaaagtaaccatttatgatattgccgaaaaactaaatatcactgcagctactgtttc | N | Hexuronate utilization operon transcriptional |
| WP_012026232.1 | 12 | 0.00037 | **5.7** | atggcagcgcagaaaaaattgataaaagttcacccaaccgacaatgtagcggttgctttggtg | N | Altronate hydrolase (EC 4.2.1.7) |
| WP_012026233.1 | 14 | 0.00003 | **7.7** | atggaaaaaattaacagatcaaattcagagttttcaaacaggcttccaattaaaattgtacaattt | N | Altronate oxidoreductase (EC 1.1.1.58) |
| WP_012026234.1 | 3 | 0.00005 | **6.8** | atggcaaaatattcaagaatagaggttgcgcagacaatgaaagataacggaatggtgccgttg | N | 4-hydroxy-2-oxoglutarate aldolase (EC 4.1.3.16) @ 2-dehydro-3- |
| WP_044048511.1 | 6 | 0.00013 | **3.8** | atgagtagagtagttgcatttggagaaatcatgctgcgtttatcgacagaaagacatttacgttttt | N | 2-dehydro-3-deoxygluconate kinase (EC 2.7.1.45) |
| WP_012026236.1 | 15 | 0.00006 | **6.4** | atgagcgcaaatacattcatacacgacaatttttattagaaaataaatacgctgaagagttatat | N | Uronate isomerase (EC 5.3.1.12) |
| WP_012026237.1 | 11 | 0.01289 | **3.5** | atgacaaacttatttgacataaaaggaaaagttgcccttattacaggaagtacgcacggactgg | N | 2-deoxy-D-gluconate 3-dehydrogenase (EC 1.1.1.125) |
| OXE95634.1 | 7 | 0.004 | **4.8** | atgacaaaatatagttcaagatacgcgtcaagtccagaagctgtaaaaaaatatgatacacaa | N | 4-deoxy-L-threo-5-hexosulose-uronate ketol-isomerase (EC 5.3.1.17) |
| WP_012026239.1 | 3 | 0.00163 | **4.6** | atgagtcaaaccgatgctgttacagtaaatcaaaccgttaagtccgcaggaagatatcgttgga | N | Hexuronate transporter |
| WP_012026240.1 | 5 | 0.73526 | **1** | ttgaaaaacacaaaaaccctttatttactcactgtcctgctttttcaggaattgtattacattcaagtg | Y | Xylanase |

Cluster 2

| NCBI annotation | # unique peptides | Pectin/DDW: P-Value | Pectin/DDW: Fold change | Gene Sequence | Signal Peptide? | RAST annotaion |
|---|---|---|---|---|---|---|
| WP_012022292.1 | 10 | 0.00148 | **203.9** | cgtaacagagattttagttcagccaggtaatataacccacaccataacagcagaagaaaaag | Y | hypothetical protein |
| WP_012022293.1 | 15 | 0.00954 | **8.4** | ttaaagcatggggtgatgttcctgctcgttttgaacctatcactacagcaactttatatttgcctaaat | Y | putative outer membrane protein, probably involved in nutrient binding |
| WP_012022294.1 | 34 | 0.00139 | **22.1** | gtaacatcagcagagggttcgcctgatgccgatatacgaattagagttcgtggaggaggatcat | Y | TonB family protein / TonB-dependent receptor |

Cluster 3

| NCBI annotation | # unique peptides | Pectin/DDW: P-Value | Pectin/DDW: Fold change | Gene Sequence | Signal Peptide? | RAST annotaion |
|---|---|---|---|---|---|---|
| WP_012022634.1 | 3 | 0.10566 | **1.8** | atgaaaatcaattttaaaaaaacgtacaaatttgctataaaatcggcattgtatatcagtcttttctc | N | Phosphate regulon sensor protein PhoR (SphS) (EC 2.7.13.3) |
| SEP16691.1 | 3 | 0.04429 | **2.6** | ttgctatttatgaaaaaaacacaaactaagattttattagttgacgacgaaccggatatcttagaa | N | Response regulator receiver:Transcriptional regulatory protein, C-terminal |
| WP_012022636.1 | 30 | 0.05111 | **3.6** | atgaacaatgatccgttgccttttgcaaatgttttagttaaaggtacaaacattagcgtaaataccg | N | TonB-dependent receptor |
| WP_012022637.1 | 22 | 0.00997 | **14.4** | atgaaaaaatcaattttaaatttcgttacagttgttgctttatcaggaggtttacttacaagctgttcta | Y | FIG00649626: hypothetical protein |
| WP_012022639.1 | 1 | 0.00753 | **47.1** | ttgattacagccagcgatatatttacaatttcaagccataaacaatttgaaaaaacagcattaaa | N | Long-chain-fatty-acid--luciferin-component ligase (EC 6.2.1.19) |

Cluster 4

| NCBI annotation | # unique peptides | Pectin/DDW: P-Value | Pectin/DDW: Fold change | Gene Sequence | Signal Peptide? | RAST annotaion |
|---|---|---|---|---|---|---|
| WP_012026068.1 | 5 | 0.02085 | **2.9** | atgaaaacaaaatttaataaatatatatcactggcacttttacttattgtaggtgcttcgtgcagcga | Y | putative outer membrane protein, probably involved in nutrient binding |
| WP_012026069.1 | 5 | 0.05904 | **5.4** | atgaaacaagcacttttaaagagatgtagttttcttttgtttgcactgatgagtgtaattacttatgctc | Y | TonB family protein / TonB-dependent receptor |
| WP_012026070.1 | | | | atgaaatacattctaaccttatttttaataacaacactttcaatatcagcccaaacgcttgataacaaattgacattaa | | rhamnogalacturonan acetylesterase |

| | | | | | | |
|---|---|---|---|---|---|---|
| WP_012026071.1 | 2 | 0.00094 | **4.4** | atgaaaaagaacatcatcatatcctctttatttctttcaaccgtaatctttgcgcagaacaaaggttt | Y | Putative glycosyl hydrolase of unknown function (DUF1680) |
| WP_012026072.1 | 7 | 0.18079 | **2.5** | atgataaagaaattaataagcggtgctgttttgtccttgcttttgaccacaaacggacaagcacaa | Y | COG3866 Pectate lyase |

*Validated by qPCR

**Table S4- Pectin induced up regulated SusC/D proteins in *F. johnsoniae***

**SusC-like protein**

| NCBI annotation | DNA seq. | fold change | p-Value | Predicted susbtrate |
|---|---|---|---|---|
| WP_012022294.1 | atgaattttaaagatttattaaacaaaggagcaa | 22.12 | 0.00139 | |
| WP_012026069.1 | atgaaacaagcacttttaaagagatgtagttttct | 5.38 | 0.05904 | pectins |
| WP_012026168.1 | atgtgtacagaccgaaaaatcaaatcgtttaaa | 102.43 | 0.00484 | |
| WP_012026229.1 | atgaacattcaaaaatcattaaagaaaaaaat | 8.39 | 0.00035 | pectins |

**SusD-like protein**

| NCBI annotation | DNA seq. | fold change | p-Value | Predicted susbtrate |
|---|---|---|---|---|
| WP_012022293.1 | atgaaacataaattaataatagcaggattgata | 8.36 | 0.00954 | |
| WP_012024115.1 | atgaaatatagttttaaaataaaaacattaggag | 11.61 | 0.03876 | |
| WP_052295191.1 | atgaaaaatataaaaataacactttcattattatt | 3.22 | 0.04623 | peptides |
| WP_012025858.1 | atgaaacgatatatttttagaaatttaataatagg | 2.46 | 0.00397 | hemicelluloses |
| WP_012026068.1 | atgaaaacaaaatttaataaatatatatcactgg | 2.87 | 0.02085 | pectins |
| SHH12854.1 | atgttaggaatgattgttacaagcttaagttcttgc | 11.86 | 0.00449 | pectins |

**TableS5- List of primers for qPCR**

| Primer name | NCBI acession number | RAST annotation | Forward primer | Reverse primer | Tm | Amplicon length |
|---|---|---|---|---|---|---|
| **Target genes** | | | | | | |
| TonB 445 | WP_012022294. | TonB family protein / TonB-dependent receptor | GGGTATAGACTGCCTCCTGTAA | TCCTCCTCCACGAACTCTAAT | 62 | 92 |
| Transcript_regul YafY fmaily | WP_012022876. | transcriptional regulator, putative | GGGTATAGACTGCCTCCTGTAA | GCGTAGTGTGTTCCCAAAGA | 63/62 | 110 |
| auxin_regulted protein | WP_012023111. | putative auxin-regulated protein | ACAGCGGGTACCAACATTGTAA | CGTGGTACGGCAGTCCATT | 58 | 79 |
| SusD_73 | SHH12854.1 | RagB/SusD domain protein | CCTGCCGATGCAACCTATAA | AACCACGGATTTCCTCCATAAA | 62 | 98 |
| TonB_260 | WP_012026229. | TonB family protein / TonB-dependent receptor | GCTTTCAAACGCAGGAAGTAAG | GAACCGTATCCTACAACCACTAC | 62 | 106 |
| TonB_2144 | WP_012026069. | TonB family protein / TonB-dependent receptor | GGCGCTGGCTTGTTCTTTAT | TGATTGTTGCATCCCATTTAATATCT | 58/59 | 132 |
| Pectate-lyase | WP_012026072. | COG3866 Pectate lyase | AACTGACGGAGGAGCAAAC | ACCTTCGTGTTGTGCATTTAAG | 62 | 100 |
| RemA_847 | WP_012022896. | internalin, putative RemA | GGAGAGACAAATAGCGGTACAA | ATGGCCTATCCAGGTGTTATTT | 62 | 102 |
| **Refrence genes** | | | | | | |
| Electron transfer flavoprotein, alpha subunit- ETF | WP_012023552. | Electron transfer flavoprotein, alpha subunit, Threonine synthase | TTTAACCCGACACTTGGAGAC | CGATATCAGCATCGGCAATAGA | 62 | 94 |
| DNA gyrase subunit B (EC 5.99.1.3) -GyrB | WP_012024321. | DNA gyrase subunit B | GAGAGGTTGTATCTCCGGTTTC | GAGCCTGAGCTGCTAAGATTAC | 62 | 117 |
| 16S rRNA | | 16S rRNA | CGGCAACGAGCGCAACCC | CCATTGTAGCACGTGTGTAGCC | 55 | 130 |

**Table S6 - Predicted operons in TonB related cluster 1**

**Tool 1-**

**Operon Mapper: Bacterial Operon Prediction**

| Operon | NCBI annotation | Sequence | gene function | start | end | Strand | Operon mapper annotation |
|---|---|---|---|---|---|---|---|
| 1 | OXE95634.1 | atgacaaaa | 4-deoxy-L-threo-5-hexosulose-uronate ketol-isomerase (EC 5.3.1.17) | 215 | 1054 | + | 5-keto 4-deoxyuronate isomerase |
| 1 | WP_012026237.1 | atgacaaac | 2-deoxy-D-gluconate 3-dehydrogenase (EC 1.1.1.125) | 1059 | 1850 | + | R] Dehydrogenases with different specificities |
| 1 | WP_012026236.1 | atgagcgca | Uronate isomerase (EC 5.3.1.12) | 1875 | 3278 | + | Glucuronate isomerase |
| 1 | WP_044048511.1 | atgagtaga | 2-dehydro-3-deoxygluconate kinase (EC 2.7.1.45) | 3432 | 4478 | + | Sugar kinases, ribokinase family |
| 1 | WP_012026234.1 | atggcaaaa | 4-hydroxy-2-oxoglutarate aldolase (EC 4.1.3.16) @ 2-dehydro-3-deoxy | 4491 | 5159 | + | 2-keto-3-deoxy-6-phosphogluconate aldolase |
| 1 | WP_012026233.1 | atggaaaaa | Altronate oxidoreductase (EC 1.1.1.58) | 5333 | 6817 | + | Mannitol-1-phosphate/altronate dehydrogenases |
| 1 | OXE95634.1 | atgacaaaa | 4-deoxy-L-threo-5-hexosulose-uronate ketol-isomerase (EC 5.3.1.17) | 6856 | 8478 | + | Altronate dehydratase |
| 2 | WP_012026231.1 | atgagcgaa | Hexuronate utilization operon transcriptional repressor ExuR | 8719 | 9747 | - | Transcriptional regulators |
| 3 | WP_012026230.1 | ttgactatatc | Polygalacturonase (EC 3.2.1.15) | 10038 | 11465 | + | Endopolygalacturonase |
| 3 | WP_012026229.1 | atgaacattc | TonB family protein / TonB-dependent receptor | 11511 | 14615 | + | |
| 3 | SHH12854.1 | atgttaggaa | RagB/SusD domain protein | 14628 | 16301 | + | |
| 4 | WP_012026227.1 | ttgattttaatg | Pectinesterase (EC 3.1.1.11) | 16406 | 17362 | + | Hydrolases of the alpha/beta superfamily |
| 4 | WP_012026226.1 | atgaactata | Rhamnogalacturonides degradation protein RhiN | 17383 | 18603 | + | Predicted unsaturated glucuronyl hydrolase |
| 4 | WP_012026225.1 | atgataaaga | Rhamnogalacturonides degradation protein RhiN | 18616 | 19821 | + | Predicted unsaturated glucuronyl hydrolase |
| 4 | WP_044048002.1 | atgtcggtttta | Rhamnogalacturonides degradation protein RhiN | 19871 | 21064 | + | Predicted unsaturated glucuronyl hydrolase |

**Tool 2-**

**FGENESB: Bacterial Operon and Gene Prediction**

| Operon | NCBI annotation | Sequence | gene function | start | end | Strand | score |
|---|---|---|---|---|---|---|---|
| 1 | OXE95634.1 | atgacaaaa | 4-deoxy-L-threo-5-hexosulose-uronate ketol-isomerase (EC 5.3.1.17) | 215 | 1054 | + | 470 |
| 1 | WP_012026237.1 | atgacaaac | 2-deoxy-D-gluconate 3-dehydrogenase (EC 1.1.1.125) | 1059 | 1850 | + | 587 |
| 1 | WP_012026236.1 | atgagcgca | Uronate isomerase (EC 5.3.1.12) | 1875 | 3278 | + | 725 |
| 2 | WP_044048511.1 | atgagtaga | 2-dehydro-3-deoxygluconate kinase (EC 2.7.1.45) | 3456 | 4478 | + | 687 |
| 2 | WP_012026234.1 | atggcaaaa | 4-hydroxy-2-oxoglutarate aldolase (EC 4.1.3.16) @ 2-dehydro-3-deoxy | 4491 | 5159 | + | 422 |
| 3 | WP_012026233.1 | atggaaaaa | Altronate oxidoreductase (EC 1.1.1.58) | 5369 | 6817 | + | 1054 |
| 3 | WP_012026233.1 | atggaaaaa | 4-deoxy-L-threo-5-hexosulose-uronate ketol-isomerase (EC 5.3.1.17) | 6856 | 8478 | + | 1196 |
| 4 | WP_012026231.1 | atgagcgaa | Hexuronate utilization operon transcriptional repressor ExuR | 8719 | 9747 | - | 651 |
| 5 | WP_012026230.1 | ttgactatatc | Polygalacturonase (EC 3.2.1.15) | 10038 | 11465 | + | 717 |
| 5 | WP_012026229.1 | atgaacattc | TonB family protein / TonB-dependent receptor | 11511 | 14615 | + | 1667 |
| 5 | SHH12854.1 | atgttaggaa | RagB/SusD domain protein | 14646 | 16301 | + | 1049 |
| 6 | WP_012026227.1 | ttgattttaatg | Pectinesterase (EC 3.1.1.11) | 16406 | 17362 | + | 335 |
| 6 | WP_012026226.1 | atgaactata | Rhamnogalacturonides degradation protein RhiN | 17383 | 18603 | + | 872 |
| 6 | WP_012026225.1 | atgataaaga | Rhamnogalacturonides degradation protein RhiN | 18616 | 19821 | + | 799 |
| 6 | WP_044048002.1 | atgtcggtttta | Rhamnogalacturonides degradation protein RhiN | 19871 | 21062 | + | 848 |

*Validated by qPCR
